# Living cell mTORC1 inhibition reporter mTIR reveals nutrient-sensing targets of histone deacetylase inhibitor

**DOI:** 10.1101/2023.05.19.541400

**Authors:** Canrong Li, Yingyi Ouyang, Chuxin Lu, Fengzhi Chen, Yuguo Yi, Shujun Peng, Yifan Wang, Xinyu Chen, Xiao Yan, Shuiming Li, Lin Feng, Xiaoduo Xie

**Affiliations:** School of medicine, Shenzhen Campus of Sun Yat-sen University, Sun Yat-sen University, Shenzhen, China; College of Life Sciences and Oceanography, Shenzhen Key Laboratory of Microbial Genetic Engineering, Shenzhen University, Shenzhen, China; Department of Experimental Research, State Key Laboratory of Oncology in South China, Collaborative Innovation Center for Cancer Medicine, Sun Yat-sen University Cancer Center, Guangzhou, China

**Author notes:** Correspondence (X.X). These authors contributed equally (Canrong Li, Yingyi Ouyang, Chuxin Lu).

**Keywords:** mTORC1, Live-cell reporter, HDAC inhibitor, Panobinostat, Amino acid sensing

## Abstract

Mammalian or mechanistic target of rapamycin complex 1 (mTORC1) is a clinically effective therapeutic target for diseases such as cancer, diabetes, aging, and neurodegeneration, yet an efficient tool to monitor mTORC1 inhibition in living cells or tissues is still lacking. Here we devised a genetically encoded mTORC1 inhibition reporter termed mTIR that exhibits a highly contrasted fluorescence puncta pattern in response to mTORC1 inhibition. mTIR specifically senses physiological, pharmacological and genetic inhibition of mTORC1 signaling in living cells and tissues. Importantly, mTIR can be applied as an powerful tool for imaging-based visual screening of mTORC1 inhibitors. By this method, we identified histone deacetylase inhibitors (HDACi) that selectively inhibit mTORC1 by inducing nutrient-sensing gene expression. Thus, mTIR is a unique living cell reporter efficiently detecting the inhibition of mTORC1 activity, and the HDACi Panobinostat transcriptionally target mTORC1 signaling via amino acids sensing.

## INTRODUCTION

The highly conserved serine/threonine protein kinase mammalian or mechanistic target of rapamycin (mTOR) plays central roles in cell metabolism by coordinating cellular and extracellular signals such as growth factor (GF), amino acid (AA) (Kim and Guan, 2019; Liu and Sabatini, 2020). Dysregulation of the mTOR signaling is associated with human diseases including cancer, diabetes, aging, and mTORopathies (Inoki et al., 2005; Saxton and Sabatini, 2017).

mTOR nucleates two structurally and functionally distinct mTOR complexes (mTORC) namely mTORC1 and mTORC2 to control cell growth by phosphorylating numerous substrates such as eukaryotic initiation factor 4E (eIF4E) binding protein 1 (4EBP1), ribosomal S6 kinase (S6K), and protein kinase B (PKB or AKT) respectively (Battaglioni et al., 2022). Decades of efforts have established mTORC1 as a master controller in cellular anabolic processes, including the synthesis of proteins, lipids, and other macromolecules, and in catabolic processes such as autophagy (Liu and Sabatini, 2020; Mossmann et al., 2018). mTORC1 is a lozenge-shaped dimer containing the mTOR kinase, the regulatory-associated protein of mTOR (Raptor), the mammalian lethal with SEC13 protein 8 (mLST8), the DEP domain-containing mTOR-interacting protein (DEPTOR), and the proline-rich Akt substrate of 40 kDa (PRAS40) (Sabatini, 2017; Saxton and Sabatini, 2017). Under nutrient-rich conditions, mTORC1 is recruited by Rag GTPases (complexed as RagA/B and RagC/D heterodimers) to the lysosomal membrane, where it is activated by the allosteric binding of RheB GTPase, with both Rag and RheB GTPases anchored to the lysosomes (Kim et al., 2008; Sancak et al., 2008). GTP loading of RheB is essential for mTORC1 activation, which is negatively regulated by the upstream GTPase-activating protein (GAP) tuberous sclerosis complex (TSC) (Inoki et al., 2003; Zhang et al., 2003). The tumor suppressor TSC (TSC1/TSC2) integrates GF and stress-sensing signaling pathways, including PI3K/AKT, LKB1/AMPK, Wnt/GSK3, and ERK/RSK to regulate the interaction between RheB and mTORC1. Gene mutations in these signaling pathways deregulate mTORC1 activity (Inoki et al., 2005; Torrence and Manning, 2018). AA-sensing is another essential mechanism for mTORC1 activation. Essential amino acids (EAA) such as leucine (Leu), arginine (Arg), and methionine (Met) have been reported to bind to their sensor proteins to promote GTP loading of Rags via inhibition of GATOR1 GAP activity (Wolfson and Sabatini, 2017). Oncogenic mutations of GF signaling genes or AA-sensing genes such as PI3K, AKT, TSC, and GATOR1 lead to hyperactivation of mTORC1 and promote uncontrolled cell growth and malignant transformation (Liu and Sabatini, 2020; Torrence and Manning, 2018). Inhibition of mTORC1 signaling has been shown to be clinically effective in human cancer therapy (Qiu et al., 2021; Thoreen et al., 2009).

Given the essential role of mTORC1 in cell growth and its high relevance to various diseases, it is important to monitor the inhibition of mTORC1 in cells and exploit new inhibitors for precision medicine. Methods to detect mTORC1 activity were mostly immunochemical techniques based on antibodies recognizing specific phosphorylation sites of mTORC1 substrates, such as phospho-4EBP1 (T37/T46) antibodies, phospho-S6K1 (T389) antibodies, and phospho-S6 (S235/S236 or S240/S244) antibodies recognizing the downstream ribosomal protein S6. However, techniques such as immunoblotting, immunofluorescence, immunohistochemistry, and fluorescence-activated cell sorting are strictly dependent on the specificity of these phosphor antibodies and require careful scaling to avoid background signals (Saper, 2009), and it is expensive and cumbersome to use antibodies for high-throughput screening purposes. In addition, antibody-based methods typically require disruption or fixation of cells or tissues under nonphysiological conditions. To overcome such limitations, fluorescent kinase reporters for mTORC1 have been developed recently. For example, TORCAR and AIMTOR are genetically encoded reporters that allow noninvasive detection of mTORC1 activation by live-cell imaging. (Bouquier et al., 2020; Zhou et al., 2015). These reporters are valuable tools to study mTORC1 kinase signaling at subcellular resolution and have recently been used to localize compartmentalized mTORC1 activity in the nucleus (Zhong et al., 2022; Zhou et al., 2020). However, the small changes in fluorescence ratio between acceptor and donor fluorophores, as well as the strong autofluorescence and light scattering effects, limit their use in drug screening and in tissues (Kardash et al., 2011). In addition, TORCAR requires quantitative imaging for a single cell, which limits its application to heterogeneous cell populations (Bouquier et al., 2020;

Zhou et al., 2015). Other kinase-dependent GFP translocation-based reporters (KTRs) such as GFP-LC3 or GFP-TFEB, which change their subcellular localization upon mTORC1 inactivation, have been flawed either for their low spatial resolution or for nonspecific responses (Regot et al., 2014). Therefore, alternative fluorescence reporters are needed to visualize mTORC1 activity in living cells and tissues. Here, we developed a live cell mTORC1 inhibition reporter (mTIR) with a simple expression and a high contrast signal pattern to specifically detect mTORC1 inhibition in cultured cells and in tissues.

Histone deacetylases (HDACs) remove acetyl groups from acetyl-lysine residues in histones and non-histone proteins, which play a central role in numerous biological processes, mainly by modulating chromatin and regulating gene transcription (Haberland et al., 2009; Li and Seto, 2016). HDACs are essential epigenetic regulators in many human tumor cells. Histone deacetylase inhibitors (HDACi) such as Panobinostat and Entinostat have been approved for the treatment of hematologic malignancies with broad target specificity via multiple mechanisms, including induction of cell death, cell cycle arrest, apoptosis, differentiation, and promotion of immunogenicity (Haberland et al., 2009; Li and Seto, 2016). Nevertheless, how HDACi affect key metabolic signaling pathways such as mTORC1 is largely unexplored. Here, mTIR was used for visual screening of compounds that inhibit mTORC1 in live cells. We found that HDACi are strongly targeting mTORC1, and a subsequent mechanistic study revealed a novel mechanism of action (MOA) of HDACi in transcriptional regulation of AA sensor genes.

## RESULTS

### Design and characterization of mTIR

To directly visualize the inactivation of mTORC1 in living cells, we developed a synthetic fluorescent biosensor called mTIR (mTORC1 inactivation reporter) inspired by SPARK and SiRI reporters (Linghu et al., 2020; Zhang et al., 2018). mTIR consists of two parts: First, HA -tagged 4EBP1 was fused to the fluorescent protein mCherry, followed by a homo-hexameric tag 3 (HOTag3); a self-cleaving P2A peptide joined the second part: Flag-eIF4E fused to the C-terminal homo-tetrameric tag 6 (HOTag6) (Figure 1A). The 17-kDa 4EBP1 is a high-quality mTORC1 substrate that binds strongly to the 25-kDa eukaryotic initiation factor 4E (eIF4E) in its dephosphorylated forms (Bohm et al., 2021; Peter et al., 2015). To initiate cap-dependent mRNA translation, activated mTORC1 hierarchically phosphorylates at least four sites: Thr37, Thr46, Ser65, and Thr70, releasing eIF4E from the tightly bound 4EBP1:eIF4E complex to subsequently form the translation initiation complex (Figure S1A) (Gingras et al., 1999; Gingras et al., 2001; Marcotrigiano et al., 1999). When mTORC1 was activated in cells cultured under normal conditions, mTIR was expressed in a diffuse pattern. However, when mTORC1 was inhibited by serum, EAA starvation, or the inhibitor rapamycin, 4EBP1 dephosphorylated and interacted with eIF4E, linking the two parts of mTIR. This protein-protein interaction (PPI), facilitated by the polymerization of HOTags in each part, resulted in the formation of a multiplex PPI that forms bright fluorescent puncta (Figure 1A, 1B, 1C), 50%-90% of the cell population exhibited puncta patterns that were easily observed and quantified using a fluorescence microscope with a 60x oil objective. In the control group, only 10%-20% of the cells had puncta (defined by > 10 detectable puncta within the cell) (Figure 1D). The response parameters of mTIR, including percentage of puncta cells, number of puncta cells, size, response time, and doses, were quantitatively measured in different cell lines using the mTOR inhibitor Torin1. Torin1 induced a high percentage of puncta cells (70%-90%) (Figure 1E, 1F), the number (10-100 per cell) and size (1-5 μm in diameter) showed no statistical difference with different mTOR inhibitions in each cell line, although the puncta size varied between cell lines (Figure 1G, 1H). mTIR responded to Torin1 with a half-maximum inhibition time (*t*1/2) of 2.16 hours and extended to 7.06 hours to EAA starvation in U2OS cells (Figure 1I, 1J); the *t*1/2 to Torin1 prolonged to 6.23 hours in 293T cells, possibly due to lower puncta resolution (Figure S1B, 1O and S1J); and it increased with Torin1 dose reduction (Figure S1C). The half-maximum inhibitory concentration (IC50) of mTIR to Torin1 was about 11.37 nM (Figure 1K), consistent with the reported inhibitory concentration of Torin1 for 4EBP1 Thr37/T46/Ser65 phosphorylation by immunoblotting (Kang et al., 2013; Thoreen et al., 2009). These results suggested that the response of mTIR was correlated with inhibition strength and the puncta microscopic resolution in each cell lines. However, the response of mTIR to mTORC1 inhibition is independent of cell confluence (Figure S1D) or mTIR expression level. Although high mTIR expression levels indeed caused a few more background puncta (number 2, 3, and 4 clones in Figure S1E), the percentage of mTIR puncta cells remained more than 5-fold changed in response to Torin1 in all clones tested (Figure S1F). We further confirmed this by inducible Tet-On mTIR, the response of mTIR remained the same when increasing expression was induced by escalating doses of doxycycline (Dox) (Figure 1L, 1M), these results ruled out the possibility of nonspecific aggregation by mTIR overexpression.

**Figure 1.**
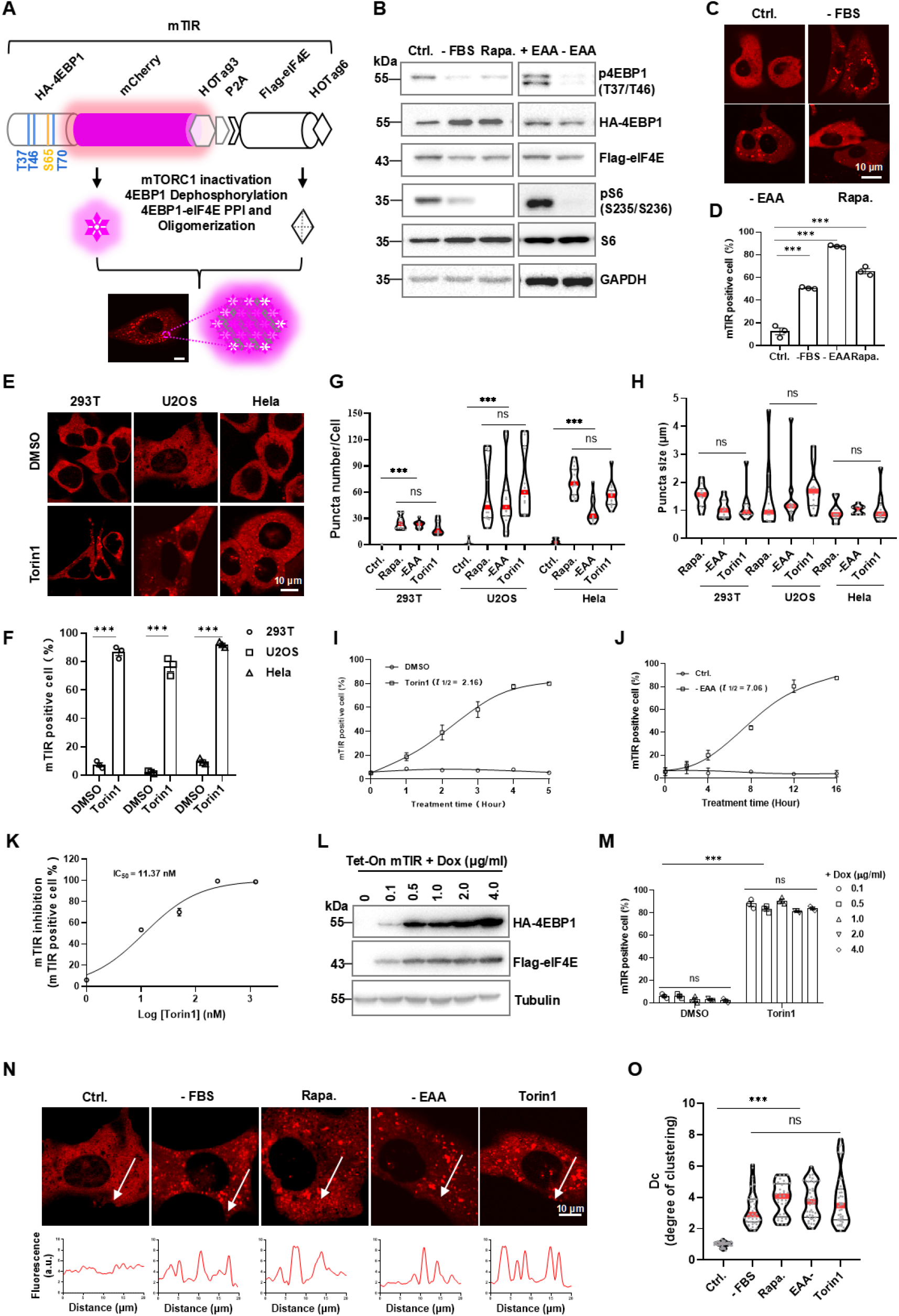
Design and characterization of mTIR. (A) Schematic diagram of the mTIR puncta formation in response to mTORC1 inactivation. (B) IB analysis of cell lysate from mTIR transfected U2OS cells treated with 100 nM rapamycin, or EAA, FBS starved for 12 hours. (C) Representative confocal Images of U2OS cells expressing mTIR as described in (B). (D) Quantification of mTIR-positive cell (defined by > 10 puncta in a cell), 3 biological replicates were photographed, about 50 cells in each sample were calculated by mTIR-positive cell percentage. (E) Representative images of mTIR in different cell lines with or without 50 nM Torin1 treatment for 12 hours. (F) Quantification of mTIR-positive cell in response to Torin1 in (E). (G) (H) Violin plots of puncta number and size in cell lines with various treatments. Red line, median; black line, interquartile range; each dot represents one cell; 15-20 cells were calculated in each sample. (I) (J) Time-response curve of mTIR for Torin1 and EAA starvation in U2OS cells. *t*1/2 were calculated with nonlinear fit analysis, 3 biological replicates at each time-point were calculated. (K) Dose-response curve of mTIR treated with various doses of Torin1 for 16 hours. IC50 was calculated with nonlinear fit analysis, 3 biological replicates at each time-point were calculated. (L) Different expression level of Tet-ON mTIR in 293T cells by dose escalating Dox induction. (M) Quantified mTIR’s responses to Torin1 with different expression level as described in (L). (N) Representative images of mTIR puncta in U2OS cells with various treatments are shown, and fluorescence histograms plotted with lines across the cells. (O) Violin plots of Dc values in U2OS cells with different treatments as described in (N). Images and quantification data are representative of at least 3 independent experiments; Error bars show standard error of mean (SEM) of 3 biological replicates for each treatment or time-point, about 50 cells in each sample were calculated by percentage; ns, no statistical significance, * P < 0.05, ** P < 0.01, *** P < 0.001, statistical analysis using two-tailed t-test; scale bar, 10 μm. See also Supplemental Figure S1.

The mTIR fluorescent puncta were easily identified in different cell lines by plotting fluorescence histograms at micrometer distances across the cells (Figure 1N, Figure S1G, S1I), and the resulting mTIR resolution could be quantitatively measured by the degree of clustering (Dc, ratio of fluorescence intensity between the puncta and the nearby cytosol) (Linghu et al., 2020). Dc values in mTORC1-inhibited cells were 2–8 times higher than in untreated cells, with higher Dc values in U2OS cells than in 293T and Hela cells (Figure 1O and S1H, S1J), which is consistent with the *t*1/2 values between cell lines (Figure 1I and S1B). Such a high-resolution mTIR signal pattern allowed us to detect the inactivation of mTORC1 easily by counting and calculating the percentages of puncta-positive cells with fluorescence images. mTIR is thus a simple and effective reporter for the inhibition of mTORC1 in living cells.

### mTIR responses to mTORC1-mediated 4EBP1 phosphorylation in living cells and in tissues

To test whether mTIR puncta formation is determined by 4EBP1 phosphorylation specifically regulated by mTORC1, phosphomimetic and non-phosphorylatable mutants of mTIR with corresponding mutations at four major mTORC1 phosphorylation sites (Thr37, Thr46, Ser65, and Thr70) in 4EBP1 were generated and then transduced into 293T cells (Figure 2A, Table S1, and Figure S2A). As expected, the mTIR^(4D)^ mutant with four mTORC1 phosphorylation sites mutated to Asp no longer responded to Torin1; however, the mTIR^(4A)^ mutant with four mTORC1 phosphorylation sites mutated to Ala continued to respond to Torin1 (Figure 2B). This could be due to the action of other sites that are directly or indirectly phosphorylated by mTORC1. To test this possibility, we generated mTIR^MT^ with all 22 Thr/Ser sites mutated to Ala except four mTORC1 sites (Figure 2A). This mutant showed the same response as the wild type to the inhibition of mTORC1 (Figure 2C, 2D). Notably, mTIR^MT(4A)^ or mTIR^MT(4D)^ in which all phosphorylation sites were mutated ablated the response, whereas mTIR^MT(4A)^ constitutively forms puncta with or without mTORC1 inhibition, and as expected, mTIR^MT(4D)^ has no puncta even when mTORC1 is inhibited in all cell lines tested (Figure 2C, Figure S2B, S2C). The discrepancy between mTIR^(4A)^ and mTIR^MT(4A)^ suggested that in addition to the four major phosphorylation sites of 4EBP1 regulated by mTORC1, there are other potential sites that affect the interaction between 4EBP1 and eIF4E. We compared mTIR with mTIR^MT^ and found that both of them responded similarly to various mTORC1 inhibitions in different cell lines, including immortalized cell lines 293T, LO2, human cancer cell lines U2OS, Hela, MCF7, A549, and mouse embryonic fibroblasts (MEF) (Figure 2D, 2E, and Figure S2D-S2F). To recapitulate the specificity of mTIR in response to 4EBP1 phosphorylation by mTORC1, we mutated the RAIP motif to AAAA (mTIR ^AAAA^), as RAIP is a major anchor site for mTORC1 to phosphorylate 4EBP1. mTIR ^AAAA^ could not be phosphorylated by mTORC1 (Beugnet et al., 2003; Tee and Proud, 2002); indeed, we observed constitutive puncta formation of mTIR ^AAAA^ mutant with or without mTORC1 inhibition, while no puncta formation was observed for the YLAA (mTIR ^YLAA^) mutant even with Torin1 treatment (Y and L mutated to A make the eIF4E binding motif “YxxxxLΦ” dysfunctional) (Mader et al., 1995) (Figure 2A, 2F and 2G). In addition, similar 4EBP1 dephosphorylation kinetics were observed for mTIR and endogenous 4EBP1 in response to Torin1 treatment (Figure 2H, 2I), suggesting mTIR authentically reflects the inhibition of cellular mTORC1 activity. These results indicated that mTIR can effectively response to mTORC1 inhibition based on the mTORC1-mediated 4EBP1 phosphorylation.

**Figure 2.**
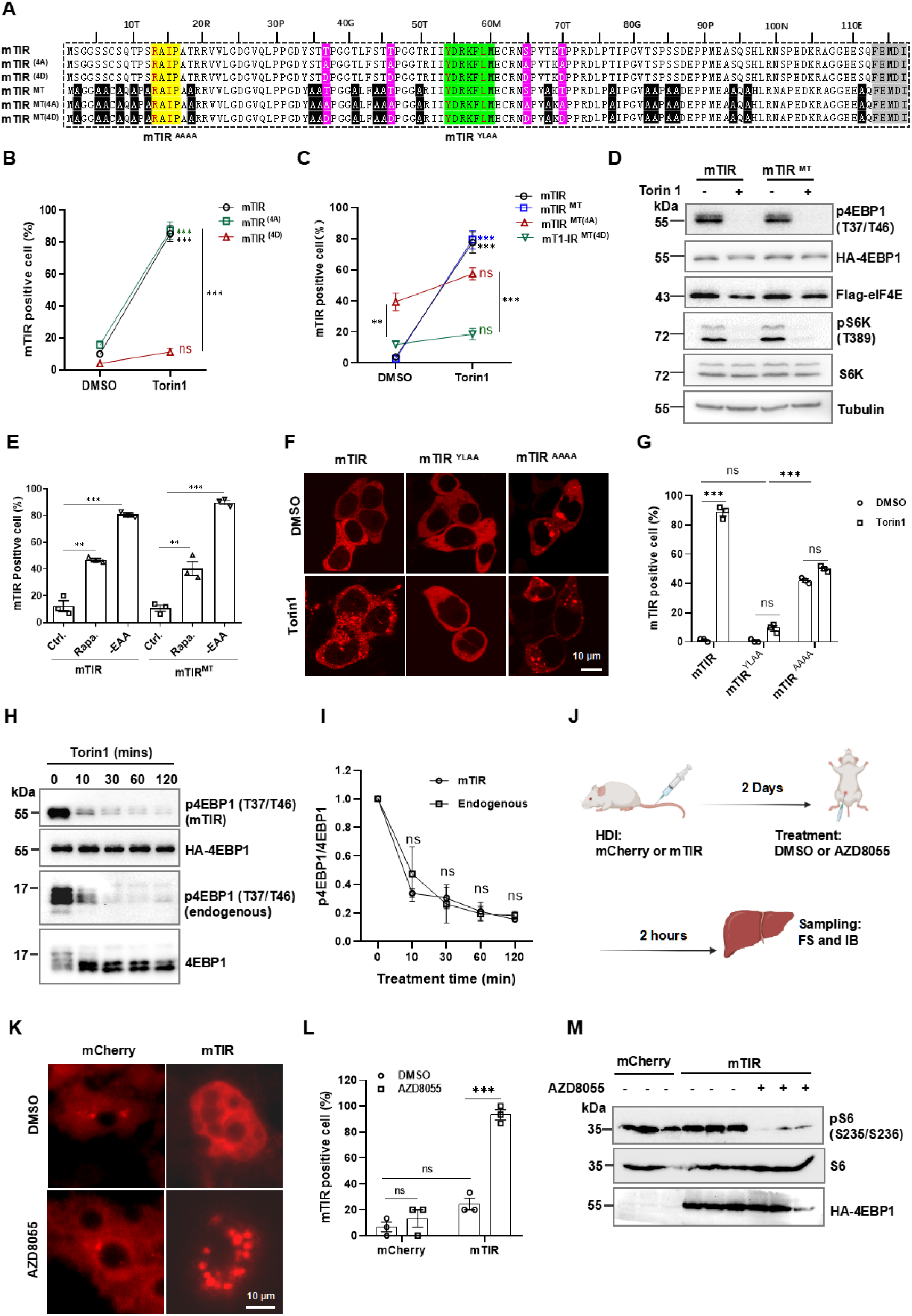
mTIR specifically responded to mTORC1-mediatd 4EBP1 phosphorylation. (A) Aligned sequences of 4EBP1 and related mutants in corresponding mTIR and mutants. (B) Responses of mTIR and mutants to Torin1 (50 nM, 12 hours) in mTIR stable-expressing 293T cells. (C) Responses of mTIR, mTIR^MT^, and mutants to Torin1 (50 nM, 12 hours) in stable 293T cell lines. (D) IB analysis of mTIR and mTIR^MT^ expressing cells in response to Torin1 (50 nM, 12 hours). (E) Responses of mTIR or mTIR^MT^ to EAA starvation or rapamycin (100 nM, 12 hours) in stable 293T cells. (F) Representative images of mTIR^AAAA^ and mTIR^YLAA^ in response to Torin1 (50 nM, 12 hours) in 293T cells. (G) Quantified responses of mTIR^AAAA^ and mTIR^YLAA^ to Torin1 in (F). (H) Representative IB analysis of 4EBP1 dephosphorylation kinetics of mTIR and endogenous 4EBP1 in response to Torin1 (50 nM, 12 hours) in 293T cells. (I) Quantified IB analysis of 4EBP1 dephosphorylation kinetics in (H), the ratio of p4EBP1/4EBP1 IB bands grayscale values were normalized to 1 at time-point 0, two independent experiments were performed. (J) Flow diagram for expressing mTIR in mouse liver tissue (HDI: hydrodynamic injection; FS: frozen sectioning, IB: immunoblotting). (K) Representative images from frozen sections of mouse liver expressing mTIR. (L) Quantification of mTIR-positive cell in frozen sections, about 50 cells were calculated in each of 3 liver samples. (M) IB analysis of liver lysate from mice tissue, details were described in the method section. Data are representative of at least 3 independent experiments; Error bar were shown with mean ± SEM of 3 biological replicates for each tested sample; ns, no statistical significance, * P < 0.05, ** P < 0.01, *** P < 0.001, statistical analysis using two-tailed t-test; scale bar, 10 μm. See also Supplemental Figure S2.

Poor resolution and high background have so far limited the application of florescence sensors such as TORCAR or AIMTOR to report mTORC1 activity in mouse tissue. To test if mTIR could report mTORC1 inhibition in tissue, we expressed it in mouse liver by hydrodynamic injection (Figure 2J). Cryosection imaging showed that mTIR expressed in a diffuse pattern and switched to intensive bright puncta in mice treated with the mTOR kinase inhibitor AZD8805 (Figure 2K-2M). These data together suggest that the puncta formation of mTIR is a dephosphorylation-specific event in response to mTORC1 inhibition both *in vitro* and *in vivo*.

### Visualization of mTORC1 signaling inhibition by inhibitors and genetic perturbation

To clarify the specificity of mTIR, we screened a compound library including 212 validated inhibitors of 103 frequently targeted protein kinases (Table S2), including those reported as mTOR dependent or independent 4EBP1 kinases such as AKT or CDK1, GSK3, ATM, *et al*. (Shahbazian et al., 2006; Shin et al., 2014; Shuda et al., 2015) (Table S3). As expected, mTIR specifically responded to inhibitors targeting PI3K/AKT/mTORC1 signaling (5 of 8 positive hits) or related upstream regulating kinases (3 of 8 positive hits) (Figure 3A, and S3A). PP121, MK-2206, GSK690693, and Buparlisib among them had already been found to be strong inhibitors of mTORC1 (referenced in Table S4), validating that our screening was effective. Other hits like GDC-0575 (CHK1 inhibitor), SKI-178 (SPHK inhibitor), CCT128930 (AKT inhibitor), and WNK464 (WNKs inhibitor) may inhibit mTORC1 through upstream signaling as reported (Table S4), which were also validated by immunoblotting assay (Figure 3B). Further exploration of the imaging data revealed that mTIR responded to none of the mTOR-independent 4EBP1 kinases summarized in Table S3 (Figure 3C); Additionally, effects of individual inhibitors targeting main putative kinases were tested by mTIR and immunoblotting side by side. As shown in Figure 3D, mTIR responded to AKT, ERK1/2 (mTOR-dependent) inhibitors, as well as different mTORC1 inhibitors with 4EBP1 dephosphorylation confirmed by IB analysis. However, we didn’t detect any response either by mTIR or by immunoblotting for those inhibitors targeting putative mTOR-independent 4EBP1 kinases such as CDK1, GSK3, ATM/ATR in normal cultured condition, possibly because those kinases might need precondition to phosphorylate 4EBP1 as previously described (Table S3). Thus, mTORC1 should be the major kinase for 4EBP1 phosphorylation in normal cultures. These results reiterated the specific response of mTIR to mTORC1-mediated 4EBP1 phosphorylation, and strongly suggested that mTIR, as it named responses to mTORC1 inhibition specifically.

**Figure 3.**
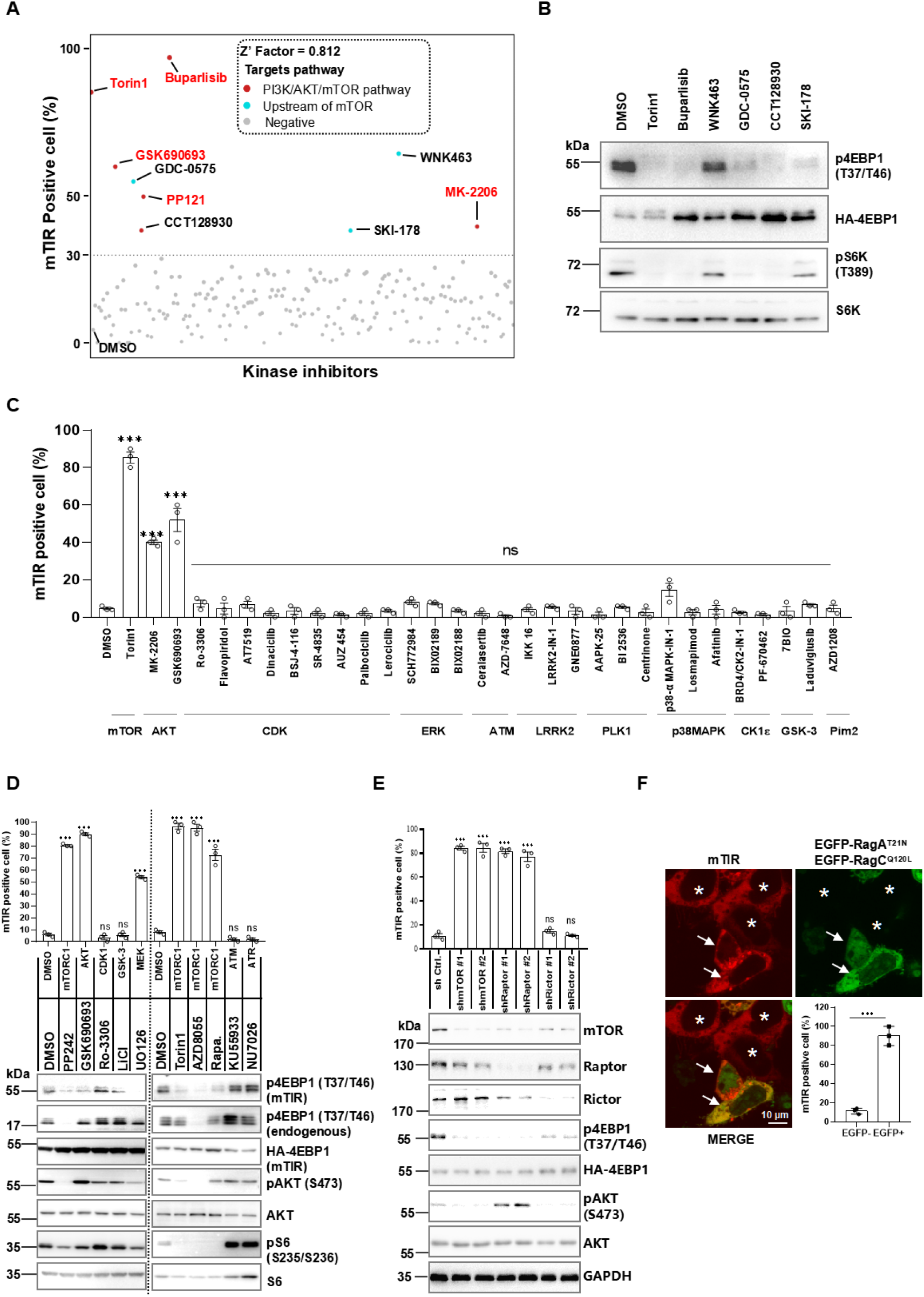
Visualizing the pharmacological and genetic inhibition of mTORC1 signaling. (A) Scatter plots of Kinase library screening by mTIR reporter, details were described in STAR methods, the red-highlighted inhibitors were found inhibiting mTORC1 in published literature, red dots denoted known PI3K/AKT/mTOR inhibitors, cyan dots denoted inhibitors possibly target mTORC1 upstream signaling as referenced in published literature summarized in Supplemental table S4, gray dots indicate no or weak inhibitory activity, the dot line shows the 30% threshold for screening. (B) IB validation of mTORC1 inhibition by hits possibly targeting mTORC1 upstream in 293T cell. (C) The responses of mTIR to inhibitors targeting mTOR-dependent and -independent 4EBP1 kinases. (D) The effects of inhibitors targeting mTOR-dependent or -independent 4EBP1 kinases by mTIR and IB analysis, 293T cells were treated for 12 hours with 100 nM PP242, 300 nM GSK690693, 1.0 μM Ro-3306, 10 mM LiCl, 10 μM U0126, 50 nM Torin1, 100 nM AZD8055, 100 nM rapamycin, 2 μM KU55933, and 10 μM NU7026 respectively. (E) The effects of mTORC1 or mTORC2 component gene knockdown on responses of mTIR in 293T cell. (F) The effects of GFP-RagA^T21N^ and GFP-RagCQ^120L^ ectopic expression on responses of mTIR in 293T cell, representative images with arrows pointing co-transfected cells, asterisks denote non-Rags expressed cell, mTIR-positive cell were quantified in GFP- and GFP+ cells. Data are representative of at least 3 independent experiments; Error bar were shown with mean ± SEM of 3 biological replicates for each tested sample; ns, no statistical significance, * P < 0.05, ** P < 0.01, *** P < 0.001, statistical analysis using two-tailed t-test; scale bar, 10 μm. See also Supplemental Figure S3, Supplemental table S2-S4.

TORCAR is a FRET-based mTORC1 sensor that effectively reports mTORC1 activity change upon GF stimulation by single cell imaging detecting FRET signal change (Zhou et al., 2015) (Figure S3B); however, no FRET signal changes was detected in the cell population of a 96-well plate neither by insulin stimulation nor by Torin1 inhibition (Figure S3C), suggesting this reporter could not be applicable to high-throughput screening by plate reading mode. In contrast, a notable Z’ factor of 0.812 indicated an excellent assay performance for mTIR in the high-throughput kinase screening (Figure 3A). When further compared to the TORCAR reporter, mTIR showed a much stronger response (2-4-fold change of Dc values) in response to Torin1 treatment compared to the little response of TORCAR by quantified single cell imaging (Zhou et al., 2016) (Figure S3D, S3E). Therefore, mTIR instead of TORCAR could be effectively applied as a high-throughput screening tool.

We further explored if mTIR could specifically report mTORC1 signaling inhibition by genetic manipulation. shRNA knockdown data verified that mTIR specifically responded to mTORC1 deficiency by Raptor or mTOR depletion but not to mTORC2 deficiency by Rictor depletion (Figure 3E). GAP protein GATOR1 inhibits Rag GTPases in responses to EAA starvation; sgRNA targeting DEPDC5, one of the subunits of GATOR1, blocked 4EBP1 dephosphorylation and the responses of mTIR induced by EAA starvation (Figure S3F-S3H). Furthermore, mTIR responded to ectopic co-expression of dominant-negative RagA^T21N^ and RagC^Q120L^ GTPases by blocking mTORC1 lysosomal recruitment (Kim et al., 2008; Sancak et al., 2008) (Figure 3F). These results indicated that mTIR is a specific tool for visualizing genetic inhibition as well as pharmacological and physiological inhibition of mTORC1 signaling.

### mTIR is reversibly modulated by mTORC1 kinase and protein phosphatase

The interaction between 4EBP1 and eIF4E is determined by 4EBP1 phosphorylation status, which is regulated by mTORC1 kinase and the protein phosphatase (Lukhele et al., 2013; Peter et al., 2015) (Figure 4A). We speculated that phosphatase may be a requisite for mTIR response; therefore, okadaic acid (OA) was used for inhibition of PP1/PP2A, two reported 4EBP1 phosphatases (Gardner et al., 2015; Liu et al., 2013). Indeed, OA efficiently blocked the puncta formation induced by AZD8055 and rapamycin (Figure 4B, 4C), and its effects on 4EBP1 phosphorylation were confirmed with biochemical analysis (Figure 4D). These results verified that 4EBP1 dephosphorylation by phosphatases was required for the response of mTIR to mTORC1 inhibition. The phosphatases also affect mTIR puncta diffusion as AA re-activation of mTORC1 cannot immediately disperse all of the mTIR puncta induced by EAA starvation, it took about 1-3 hours in 293T cells (Figure S4A, S4B), only about 10% of the cells diffused within 1 hour, 30% diffused within 3 hours, and 70% of puncta cells persisted for more than 3 hours (Figure 4E). Such slow reversibility could be attributed to the biophysical properties of mTIR puncta, which could not fully recover from photobleaching in FRAP experiments (Figure S4C, S4D, and Supplemental Video 1), and the puncta were also insensitive to the protein liquid droplet disperser 1,6-Hexanediol (HEX) (Figure S4E), suggesting mTIR did not behave as SPARK reporters (Zhang et al., 2018). To further test if phosphatases played a role in mTIR puncta diffusing processes, OA and CdCl2 were used as inhibitors for phosphatases PP1/PP2A and phosphatase PPM1G, respectively (Gardner et al., 2015; Liu et al., 2013). Indeed, OA treatment promoted up to 75% of the puncta cells to diffuse within 3 hours, despite the fact that 25% of them remained puncta-positive. (Figure 4E, S4F, S4G, and Supplemental Video 2-3). OA also prevented puncta formation induced by GSK690693 and promoted puncta diffusion after drug washing out. The PPM1G inhibitor CdCl2 alone did not inhibit puncta formation; however, it did promote puncta diffusion (Figure 4F and 4G), suggesting different types of phosphatases may act differently on the response of mTIR. These results overall suggested that mTIR was reversibly regulated by mTORC1 and phosphatases with slow kinetics, which would be advantageous to record a transient inhibition of mTORC1 since the puncta were biophysically stable upon mTORC1 re-activation.

**Figure 4.**
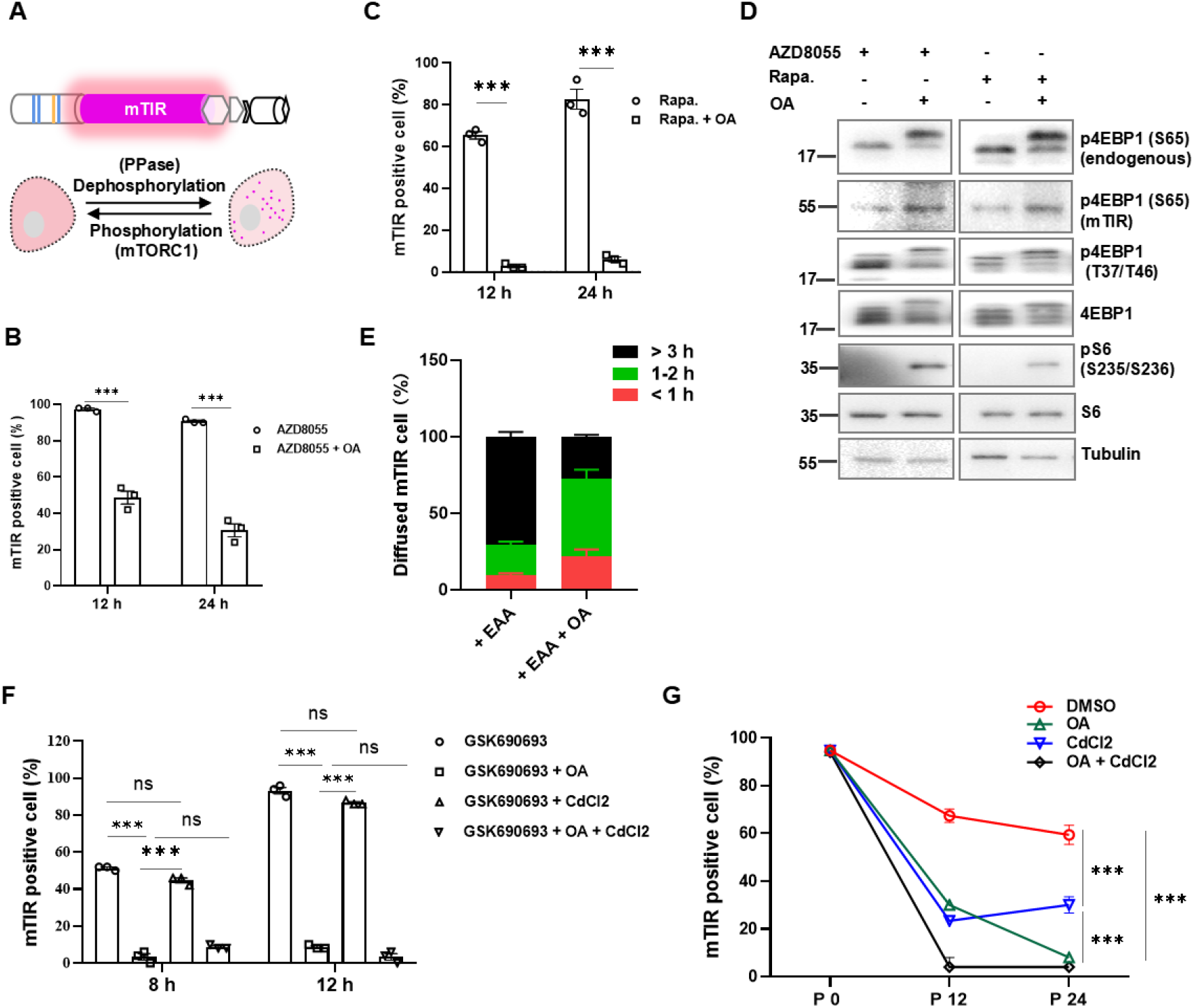
mTIR is modulated by mTORC1 kinase and protein phosphatases. (A) Diagram of mTIR regulation by mTORC1 and protein phosphatases. (B) Effect of OA on mTIR puncta formation induced by AZD8055 in 293T cells. (C) Effect of OA on the response of mTIR to rapamycin in 293T cells. (D) IB analysis of OA’s effect on 4EBP1 phosphorylation in response to mTORC1 inhibition by AZD8055 and rapamycin. (E) mTIR puncta diffusion analysis upon refeeding of EAA with or without OA pre-treatment in 293T cells (F) Effects of OA or CdCl2 or combined treatment on mTIR’s responses to mTORC1 inhibition by GSK690693. (G) Time course effects of OA or CdCl2 or both on the diffusion of mTIR puncta after GSK690693 washing out. 20 nM OA, 5.0 μM CdCl2, 100 nM GSK690693, 100 nM AZD8055, 100 nM rapamycin were use for treatments; Quantification data are representative of at least 3 independent experiments; Error bar were shown with mean ± SEM of 3 biological replicates for each tested sample; ns, no statistical significance, * P < 0.05, ** P < 0.01, *** P < 0.001, statistical analysis using two-tailed t-test; See also Supplemental Figure S4, Supplemental video 1-3.

### Nuclear mTIR reports nuclear mTORC1 activity in living cells

mTORC1 activity is known to be subcellularly compartmentalized in the cytosol, in nucleus, and on various membranous organelles (Li et al., 2006; Rosner and Hengstschlager, 2008). The mTIR puncta were cytosolically localized, as 4EBP1 and eIF4E were mostly situated in the cytosol. To test if mTIR could sense the mTORC1 inhibition in the nucleus, we tagged the SV40 nuclear localization signal (NLS) to each part of mTIR (Figure 5A), leading to expression of such mTIR (NmTIR) exclusively in the nucleus (Figure 5B). Notably, NmTIR diffused in the nuclei of HCT116 and MEF cells, but it exhibited a puncta pattern in the nuclei of U2OS and 293T cells under normal culture conditions without mTORC1 inhibition (see control group in Figure 5B-5E), suggesting different nuclear mTORC1 activity in different cell lines. As previously reported (Betz and Hall, 2013; Rosner and Hengstschlager, 2008), we confirmed different nuclear mTOR expression levels in these cell lines with biochemical fractionation analysis (Figure S5A, S5B), which also showed that nuclear mTORC1 can be activated by AA stimulation in MEF but not in 293T cells, despite low basal nuclear 4EBP1 phosphorylation in both cell lines. Moreover, NmTIR effectively sensed inhibition of mTORC1 activity inputted from GF and AA signals in MEF cells (Figure 5D, 5E), despite a few background puncta cells in the control group (20%–40%), which is possibly caused by relatively low mTORC1 activity in the nucleus when compared to mTIR’s responses in the cytosol (10%–20% background puncta cells) (Figure S5C, S5D). To further confirm if NmTIR specifically senses nuclear mTORC1 activity, we compared the nuclear forms of established phosphor-mutants of mTIR^MT^ (NmTIR^MT^) in MEF and 293T cells. As expected, NmTIR^MT^ responded to Torin1, while NmTIR^MT(4D)^ did not, and NmTIR^MT(4A)^ constitutively formed puncta, suggesting NmTIR responded to mTORC1 inhibition in MEF cells (Figure 5F, 5G). However, in 293T cells, NmTIR^MT^ and NmTIR^MT(4A)^ constitutively formed puncta with or without mTORC1 inhibition, while NmTIR^MT(4D)^ did not form puncta even with Torin1 inhibition, suggesting that the constitutive puncta formation of NmTIR in the 293T nucleus was caused by a lack of nuclear mTORC1 activity (Figure 5H–5I). Overall, NmTIR is an effective tool for evaluating nuclear mTORC1 inhibition in living cells.

**Figure 5.**
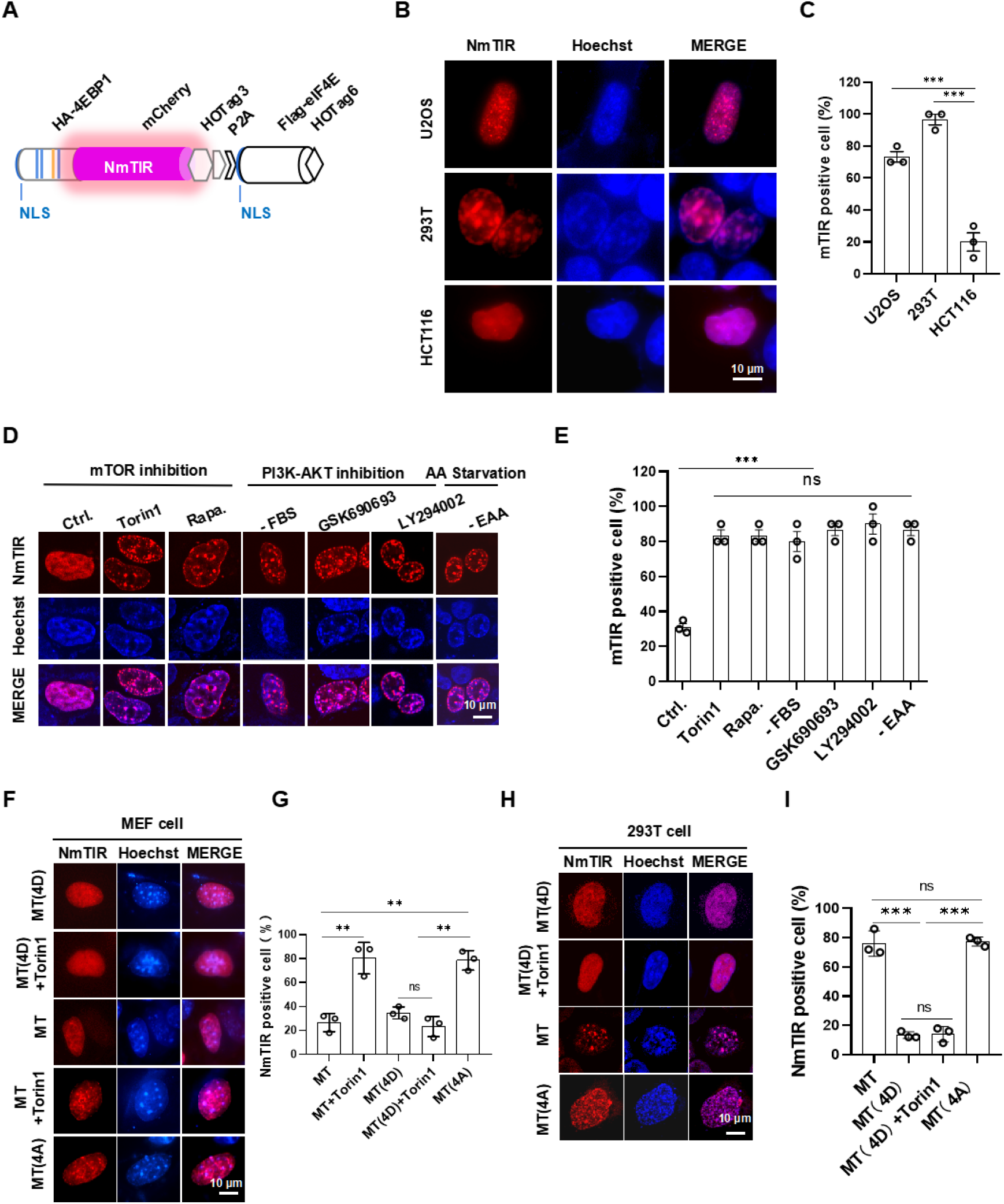
Nuclear mTIR senses mTORC1 inhibition in nucleus. (A) Structure diagram of NmTIR with NLS tags. (B) NmTIR expression pattern in the nucleus of different cell types. (C) NmTIR puncta formation in normal culture conditions in different cell types. (D) Representative images of NmTIR responding to PI3K/AKT/mTOR signaling inhibitors or EAA deprivation for 12 hours in MEF cells. (E) Quantified responses of NmTIR to PI3K/AKT/mTOR signaling inhibitors and EAA deprivation in MEF cells as in (D). (F) Representative images of NmTIR^MT^ and related mutants responding to Torin1 in MEF cells. (G) Quantified responses of NmTIR^MT^ and related mutants to Torin1 in MEF cells. (H) Representative images of NmTIR^MT^ and related mutants responding to Torin1 in 293T cells. (I) Quantified responses of NmTIR^MT^ and related mutants to Torin1 in 293T cells. Treatments: 10 μM LY294002, 50 nM Torin1, 100 nM GSK690693, 100 nM rapamycin. Images and quantification data are representative of at least 3 independent experiments; data were shown with mean±SEM of 3 biological replicates for each sample, about 50 cells in each sample were calculated by percentage; ns, no statistical significance, * P < 0.05, ** P < 0.01, *** P < 0.001, statistical analysis using two-tailed t-test; scale bar, 10 μm. See also Supplemental Figure S5.

### mTIR live-cell screening identified HDACi as mTORC1 inhibitors

mTORC1-specific inhibitors are therapeutically promising in targeted therapy of cancer and other mTOR-related diseases (Rodrik-Outmezguine et al., 2016; Xu et al., 2020). Using mTIR-expressing 293T cells, we visually screened a drug library containing 917 FDA-approved drugs and 1059 anti-tumor natural ingredients (Figure 6A). We found several clusters of drugs, including DNA damage agents, Ƴ-secretase inhibitors, and histone deacetylase inhibitors (HDACi), that had an obvious puncta-promoting effect on mTIR (Figure 6B, Figure S6A, S6B, Table S5). Most hits were DNA damage agents that inhibit DNA topoisomerases; these compounds were known to inhibit mTORC1 signaling through the p53-AMPK-TSC pathway (Ma et al., 2018); other hits, such as Rottlerin and Oridonin, had been reported as mTORC1 signaling inhibitors (Balgi et al., 2009; Daveri et al., 2016; Wang et al., 2014), which validated the effectiveness of this screening (Table S5). Positive hits were further selectively validated by immunoblotting (Figure 6C), and the most potent hits, HDACi Panobinostat and Entinostat, significantly blocked the mTORC1 activity in MCF7 (Figure 6D) and in 293T cells (Figure S6C) at 10 μM. Prominent inhibition of nuclear mTORC1 by HDACi in MEF cells were also observed with NmTIR reporter (Figure 6E, 6F). These results, together with the kinase inhibitor screening (Figure 3A), indicated that mTIR is an applicable tool for high-throughput screening of mTORC1 inhibitors in living cells.

**Figure 6.**
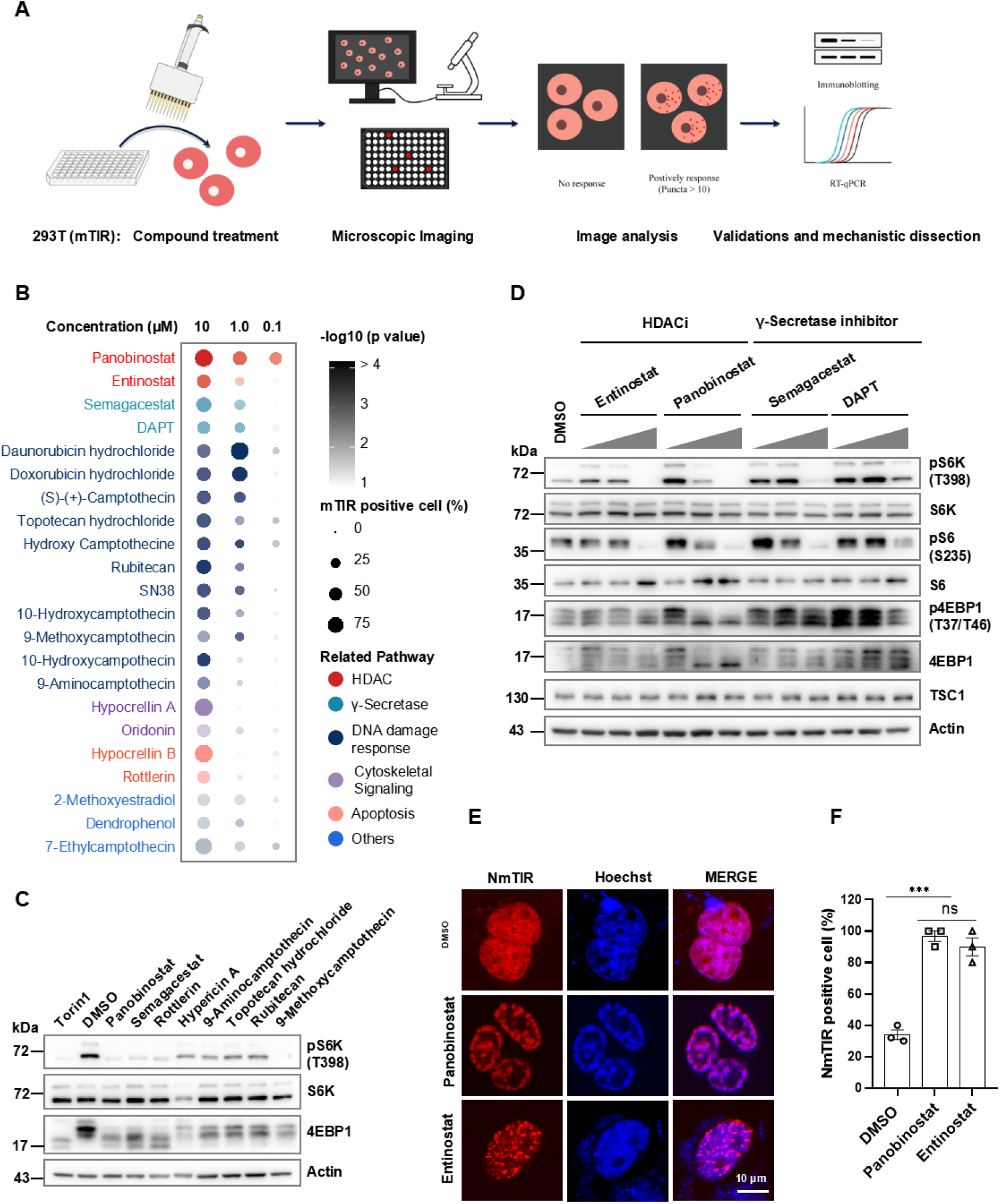
Applying mTIR to visual screening of mTORC1 inhibitors in living cells. (A) Flow diagram for live-cell screening of 1976-compound library with 293T-mTIR cell line, details were described in the methods section. (B) Bubble plot of dose-dependent responses of mTIR to screened positive hits in 293T cells. (C) IB validation of mTORC1 activity in response to the positive hits (10 μM, 12 hours) in 293T cells. (D) IB analysis of mTORC1 signaling with dose-escalation (0.1, 1.0, and 10 μM for 12 hours) treatment of HDACi and γ-Secretase inhibitors in 293T cells. (E) Representative image of NmTIR in response to HDACi (10 μM) in MEF cells. (F) Quantified responses of NmTIR to HDACi in MEF cells in (E). Images and quantification data are representative of at least 3 independent experiments; Error bars show standard error of mean (SEM) of 3 biological replicates for each treatment, about 50 cells in each sample were calculated by percentage; ns, no statistical significance, * P < 0.05, ** P < 0.01, *** P < 0.001, statistical analysis using two-tailed t-test; scale bar, 10 μm. See also Supplemental Figure S6, Supplemental table S5.

To determine if other HDACi have general inhibitory effects on mTORC1, we tested seven HDACi widely used in clinic and preclinical trials, including SAHA, TSA, Panobinostat, VPA, Chidamide, Romidepsin, and Entinostat, in various cell lines. Consistently, mTORC1 was generally inhibited in a cell-type-specific manner. For example, pan HDACi Panobinostat inhibited mTORC1 in all 3 tested cell lines, SAHA and Romidepsin had no effect on 293T cells, while VPA had little effect in MCF7 and Hela cells (Figure S6D, S6E), suggesting HDACi may exert cell-type-specific mTORC1 inhibition.

### Panobinostat inhibits mTORC1 but not mTORC2 by inducing AA sensor gene expression

Several studies reported that some HDACi, such as TSA and SAHA, could inhibit mTORC1 by targeting AKT/TSC signaling (Yan et al., 2018; Yang et al., 2020); however, neither TSC1 expression nor AKT (S473) phosphorylation changes were evident in our experiments (Yan et al., 2018; Yang et al., 2020); however, neither TSC1 expression nor AKT (S473) phosphorylation changes were evident in our experiments (Figure 6D and Figure S6C). mTORC1 was inhibited within 1 hour and reached peak inhibition after 4 hours of panobinostat treatment with no change in AKT (S473) phosphorylation (Figure S7A, S7B). We hypothesized that HDACi could suppress mTORC1 through AA signaling rather than GF signaling mediated through AKT/TSC. Indeed, Panobinostat or Entinostat almost completely inhibited AA-induced mTORC1 activation but not insulin-stimulated mTORC1 activation in 293T cells (Figure 7A, Figure S7C). Importantly, HDACi lost its inhibitory effect on mTORC1 in GATOR1-deficient cells upon AA stimulation (Figure 7B, Figure S7D), and both HDACi blocked the mTOR translocation to lysosomes upon AA stimulation, as indicated by the loss of co-localization between mTOR and LAMP2 (Figure 7C, 7D). Additionally, mTORC1 inhibition by HDACi induced dramatic autophagy after 4 hours as indicated by LC3 puncta formation, a widely reported effect of HDACi (Liu et al., 2010; Son et al., 2021) (Figure S7E, S7F). The clinical benefits of conventional mTOR inhibitors are limited by the relief of feedback inhibition of receptor tyrosine kinase (RTK) signaling, which intervenes in mTORC2 signaling (Chandarlapaty et al., 2011; Yi et al., 2020), notably, these two HDACi had no obvious effect on mTORC2 activity indicated by AKT (S473) phosphorylation, suggesting HDACi may not interfere with mTORC2 activity in those cell lines.

**Figure 7.**
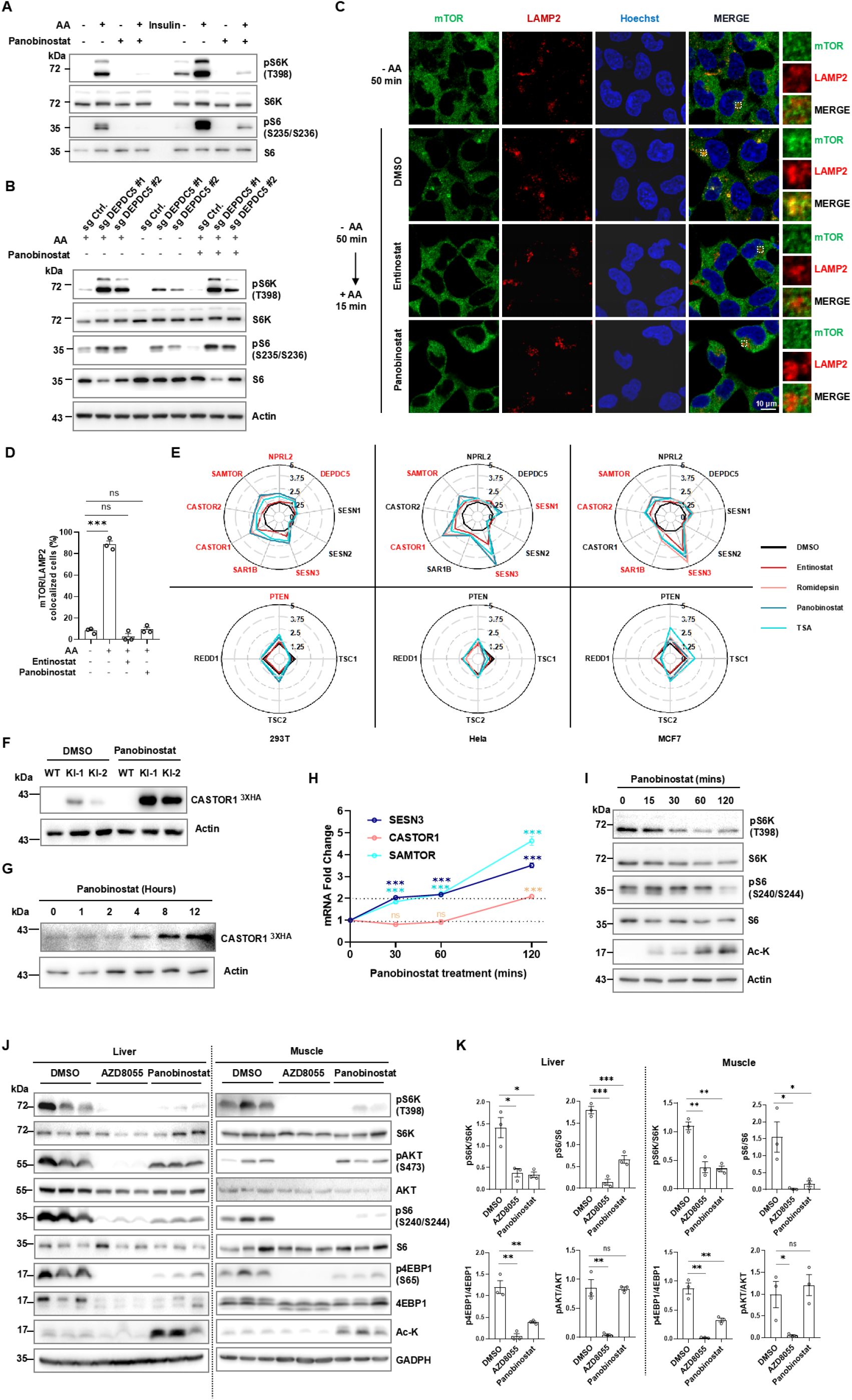
Panobinostat inhibits mTORC1 signaling by inducing AA sensor gene expression. (A) IB analysis of mTORC1 activity in response to Panobinostat with AA or insulin stimulation, 293T cells were pre-treated with DMSO or 10 μM Panobinostat for 12 hours, then starved with AA for 50 minutes (min) and stimulated with AA for 15 min, or serum starved for 12 hours and stimulated with 100 nM insulin for 15 min. (B) IB analysis of mTORC1 activity with Panobinostat treatment in DEPDC5-depleted 293T cells with or without AA stimulation. (C) Colocalization analysis by IF staining with mTOR and LAMP2 antibodies, 293T cells were pre-treated with HDACi for 12 h, then starved and stimulated with AA as indicated. (D) Percentage quantification of cells with mTOR and LAMP2 colocalization. (E) Radar plots of mRNA levels in response to HDACi treatments, relative mRNA levels of major mTORC1-inhibitory genes involved in the AA pathway and the GF pathway were quantified using RT-qPCR in 293T, Hela, and MCF7 cells treated with 4 types of HDACi for 12 h. (F) Induction of endogenously tagged CASTOR1 by Panobinostat (10 μM, 12 hours) in 293T. (G) Time course induction of endogenously tagged CASTOR1 by Panobinostat (10 μM) in 293T. (H) Time course induction of AA sensors mRNA level by Panobinostat (10 μM) in 293T. (I) Time course mTORC1 inhibition by Panobinostat (10 μM) in 293T. (J) IB analysis of mTOR signaling by Panobinostat in mouse liver and muscle tissues as described in animal methods section, samples from 3 mice per group for each treatment were showed. (K) Quantified mTOR activity (pS6K, pS6, p4EBP1, and pAKT were normalized to total S6K, S6, 4EBP1 and AKT protein levels) inhibited by Panobinostat and Torin1 in mice liver and muscle tissues in (J), all normalized values were then normalized to DMSO-treated control levels. Images and quantification data are representative of at least 3 independent experiments except those specified in figure legends; Error bars show standard error of mean (SEM) of 3 biological replicates for each treatment or time-point, about 50 cells in each sample were calculated by percentage in image data; ns, no statistical significance, * P < 0.05, ** P < 0.01, *** P < 0.001, statistical analysis using two-tailed t-test; scale bar, 10 μm. See also supplemental Figure S7.

HDACs are key epigenetic regulators that modulate numerous genes expression in cells. To explore the underlying MOA of HDACi on mTORC1 inhibition, we reviewed two RNA-sequencing data sets from the GEO database (Mackmull et al., 2015; Rampazzo et al., 2022). It revealed that major sensor genes in AA signaling, such as *CASTOR*, *SAMTOR,* and *SESTRIN,* but not the genes such as *PTEN*, *TSC1,* or *REDD1* in GF signaling, were consistently upregulated in TSA-treated glioblastoma primary cells and in Hela cells (Figure S7G). To validate this, we treated three cell lines with four HDACi and quantified the change of mRNA level for major mTORC1 inhibitory genes, including AA sensors and GATOR1 subunit genes in AA-sensing signaling, and four tumor suppressor genes downstream of GF and stress signaling, including *TSC1, TSC2, REDD1,* and *PTEN* (Figure 7E). Consistently, HDACi significantly increased the transcripts of AA sensor genes including *CASTOR1* (*CASTOR2* in MCF7), *SAMTOR,* and *SESTRIN3* in all cell lines (at least 2 fold change induced by 3 of 4 HDACi); *SESTRIN3* mRNA level was dramatically induced as previously reported (Chen et al., 2010; Zhang et al., 2015a); In contrast, only *PTEN* mRNA levels were significantly induced by HDACi treatment in 293T cells, whereas *TSC1* or *TSC2* mRNA levels exhibited negligible change in all conditions tested, which is consistent with immunoblotting results (TSC1 in Figure 6D and Figure S6C). Accordingly, the activation of mTORC1 by single amino acid (sAA) Leu, Arg, and Met (through sensors SESTRIN3, CASTOR1, and SAMTOR, respectively) was completely blocked by Panobinostat (Figure S7H). We further verified the 3xHA endogenously tagged CASTOR1 protein induction by Panobinostat (Figure 7F, 7G, Figure S7I), and observed that mRNA induction of *SAMTOR* and *SESTRIN3* was evident at 30 minutes, preceding *CASTOR1* that took 2 hours, indicating those sensor genes were induced sequentially (Figure 7H), which was well correlated with histone acetylation starting at 15 minutes and the following mTORC1 inhibition with 30-minute treatment of Panobinostat (Figure 7I). It should be noted that Panobinostat inhibits mTORC1 via the combined effects of multiple AA sensors, as single knockout of either *CASTOR1* or *SAMTOR* did not abolish the inhibition, while double knockout of both sensors obviously blunted the inhibition effect (Figure S7J-S7M). Finally, we tested the physiological effect of Panobinostat in mice. Compared to the kinase inhibitor AZD8055, Panobinostat exerted mTORC1-specific inhibition without interfering with mTORC2 activity indicated by AKT (S473) phosphorylation in liver and muscle tissues as it did in cell lines (Figure S6C, S7A, and Figure 7J, 7K). These data strongly suggest that the FDA-approved drug Panobinostat specifically targets mTORC1 without interfering with mTORC2 activity through transcriptional induction of AA sensors.

In summary, as diagrammed in Figure S7M, our study not only developed a novel genetically encoded cell reporter for mTORC1, but also revealed the MOA of HDACi in inhibiting mTORC1 signaling, which is frequently targeted in therapies for human disease.

## DISCUSSION

Basal-level mTORC1 activity is essential for maintaining cellular metabolisms in normal feeding or culture conditions, while it is inhibited when organisms are exposed to internal and external stress such as nutrient starvation, energy depletion, and genotoxic stress. This restriction of mTORC1 activity is critical for cells to keep themselves in check autonomously in response to environmental clues, preventing uncontrolled cell proliferation and other pathological damages. Therefore, an inhibition reporter for mTORC1 in living cells could expand the toolbox to facilitate research in mTOR biology.

mTIR showed a simple and highly contrasted puncta pattern upon mTORC1 inhibition in the physiological and pathological context of living cells and in mouse liver tissues, and it possesses several advantages that make it a unique mTORC1 sensor. First, it can effectively report the inhibition of mTORC1 signaling by all means, including physiological stresses (Figure 1B), pharmacological inhibition (Figure 1G and 5D), or genetic perturbation of mTORC1 signaling in living cells (Figure 3E and Figure S3D). mTIR itself did not intervene in mTORC1 signaling and exhibited no obvious toxicity in cells. Secondly, mTIR can be used as an effective tool for visual screening of mTORC1 inhibitors by imaging a heterogeneous cell population, which is infeasible for a FRET-based TORCAR reporter relaying on single cell analysis (Figure 3B, Figure S3C). In addition, NmTIR can be used as a convenient tool for monitoring mTORC1 inactivation in the nucleus (Figure 5), and importantly, mTIR effectively responded to mTORC1 inhibition in mouse liver tissues, taking advantage of its high Dc (signal/noise ratio) and long diffusing time (Figure 2J-2M).

By applying the mTIR reporter to a visual screening in living cells, we successfully identified dozens of drugs that exhibited obvious mTORC1 inhibition effect. HDACi, Panobinostat among them, was validated extensively as strong mTORC1 specific inhibitors in cell lines and in mouse tissue, mechanistically, Panobinostat potently induced nutrient sensor gene expression such as *CASTOR*, *SAMTOR,* and *SESTRIN3* by epigenetic regulation of histone acetylation (Figure 7E-7I). There were studies reporting that some HDACi inhibit mTORC1 by targeting GF signaling either by AKT (S473) phosphorylation inhibition or by TSC1 induction (Yan et al., 2018; Zhang et al., 2015b); however, none of the HDACi treatments led to evident AKT inhibition or TSC1 protein change in our experiments (Figure 7J, 7K, and Figure S7A - S7C). Whether HDACi inhibit mTORC1 through distinct signaling in a dose-dependent or tissue-specific manner needs detailed verification in animal models and in HDACi-treated clinic samples. Overall, our data are inspiring for further mechanistic dissection and clinic evaluation of HDACi in the therapy of mTOR-related diseases.

## Limitations of the study

mTIR provides limited spatial and temporal resolution in detecting mTORC1 inhibition, and its relatively slow dynamics of response makes it hard to detect transient mTORC1 activation quickly. Spatially, it can detect nuclear mTORC1 activities, but it’s hard to detect mTORC1 inhibition with mTIR on other membrane-bound organelles. Additionally, mTIR puncta cells are not discernible by FACS sorting due to small puncta size, which makes it unsuitable for Pulse Shape Analysis (Ramdzan et al., 2012). Further optimization is needed for its application to high-throughput genetic screening. Finally, validation of mTIR in transgenic animals would be crucial for further application *in vivo*.

## Acknowledgments

We thank the Core Facilities at State Key Laboratory of Oncology in South China for assistance in imaging experiments; 917 FDA approved drugs (# L1021, APExBIO) were kindly provided by professor Xinjian Liu, Sun Yat-sen University.

## Author contributions

X. Xie conceived the idea; C. Li, Y. Ouyang, C. Lu, L. Feng and X. Xie designed and performed the experiments with contributions from F. Chen, Y. Yi, Y. Wang, S. Peng, X. Chen, S. Li, and X. Yan; X. Xie, C. Li, and Y. Ouyang analyzed the data and wrote the manuscript with input from all other authors, X. Xie supervised the project.

## Data availability

All the data supporting the findings of this study are available upon request as described in STAR methods.

## STAR METHODS (for initial submission)

### RESOURCE AVAILABILITY

#### Lead Contact

Further information and requests for reagents contact Xiaoduo Xie

#### Materials Availability

All unique reagents generated in this study are available from the Lead Contact with a completed Materials Transfer Agreement.

#### Data and Code Availability

This study did not generate any unique datasets or code.

### EXPERIMENTAL MODELS

#### Cell lines

HEK293T (293T), Hela, U2OS, HCT116, LO2, MCF7, A549, and MEF cells were from ATCC and cultured in DMEM medium with 10% FBS, 100 μg/ml streptomycin, and 100 U/ml penicillin. All cells were cultured at 37°C with 5% CO2. Cells were transfected with Polyethylenimine (PEI) or Lipofectamine 3000 according to the manufacturer’s protocols. For serum starvation, cells were washed twice with PBS and cultured for 12 hours in DMEM without FBS. For EAA starvation, cells were cultured in custom-ordered AA-free DMEM with NEAA overnight, and then EAA was added for stimulation. For whole-AA starvation, cells were cultured in AA free DMEM for 50 minutes, then an EAA and NEAA mixture was added for stimulation. All cell lines were examined for mycoplasma-free status prior to experiments. Cell culture reagents used in this study were listed in the Key resources table.

#### Animal work

Mice were maintained in a pathogen-free environment. For hydrodynamic injection, 6- to 8-week-old BALB/c mice (GemPharmatech Co., Ltd.) were used for hydrodynamic tail-vein injections. 40 μg pmCherry or pmCherry-mTIR plasmids were diluted in physiological solution (0.9% NaCl) with a volume equivalent to 10% of body weight (0.1 ml/g). The whole volume was intravenously injected quickly, within 5-8 seconds. After 2 days of recovery, mice were fasted overnight and intraperitoneally injected with vehicle or AZD8055 (10 mg/kg/per mouse). 2 hours later, mice were euthanized by cervical dislocation, and liver tissues were quickly dissected out and stored at -80°C. The fluorescence images were taken with 5 μm liver cryosections prepared according to standard protocol. To test the effects of AZD8055 or panobinostat on mTOR signaling in vivo, grouped six-week old male C57 mice (GemPharmatech Co., Ltd.) (3–4 mice in each group) were injected intraperitoneally with DMSO, Panobinostat (10 mg/kg), or AZD8055 (10 mg/kg) in 30% Captisol. 8 hours post-injection, mice were euthanized and tissues were collected. Liver and muscle tissues were homogenized and lysed with RIPA buffer, total protein was quantified with BCA reagent, and immunoblotting was performed with antibodies as indicated. All animal experiments were performed following the ethical guidelines and protocols approved by the Institutional Animal Care and Use Committee (IACUC) of Sun Yat-sen University.

### METHOD DETAILS

#### Molecular cloning and plasmid constructs

Nucleic acid fragments of mTIR or TORCAR were synthesized by Tsingke Biotechnology, China, and subcloned into the pmCherry vector. All mutants were generated by standard site-directed mutagenesis method; mTIR and related mutants were also subcloned into the pCDH lentiviral vector. Tet-On mTIR were generated by subcloning mTIR into pCW57.1-teton. NmTIR and related mutants were generated by fusing SV40 NLS to the N terminals of 4EBP1 and eIF4E separately. For gene knockdown or knockout, pLKO.1-shmTOR, shRaptor, and shRictor plasmids were described in (Xie et al., 2011), and sgRNA oligos were ligated into lentiCRISPRv2 by standard protocol. All subcloning plasmids were confirmed by Sanger sequencing. Oligos for subcloning and mutagenesis were listed in Supplemental Table S6.

#### Lentivirus generation, shRNA knockdown, CRISPR/Cas9 gene knockout, and 3xHA endogenous tagging

pLKO.1 lentiviral shRNA virus and pCDH-mTIR lentiviral virus packaging and subsequent generation of stable cell lines by infection were performed as previously described in (Xie et al., 2018). Briefly, 6 well plate of 293T cells were transfected with VSVG, psPAX2, and pLKO.1 shRNA plasmids with PEI, 48 hours post-transfection, the supernatant containing virus was collected and passed through a 0.45 μm filter, and the virus was stored at −80°C until use. For lentiviral transduction, cells were infected overnight with 8 μg/mL polybrene in virus-containing medium, and cells were selected with puromycin for 2 days (48 hours) post-infection. For CRISPR/Cas9 gene knockout, individual sgRNAs were subcloned into LentiCRISPRv2 at the BsmBI site as described in the standard protocol. 293T cells were transfected with 2 µg of sgRNA plasmids for each 35-mm dish. 24 hours after transfection, 1 ug/ml of puromycin was added to refreshed medium for 2 days, and puromycin-resistant cells were pooled and amplified for IB analysis. Single cell clones were isolated using limiting dilution in 96-well plates; For sequencing, genomic DNA was extracted from the clonal cells with lysis buffer (50 mM Tris, pH 8.0, 1 mM EDTA, 0.5% Tween 20, proteinase k >0.6 U/ml), amplification of the edited sequence from genomic DNA was performed using Taq PCR mix. Purified PCR products were sequenced to identify the correctly edited clones. For 3xHA endogenous tagging of CASTOR1 in 293T cells, the sgRNA and donor sequences were designed by the online tool TrueDesign Genome (https://apps.thermofisher.cn/apps/genome-editing-portal/#/summary), constructs generation, gene targeting, and 3xHA-tagged CASTOR1 clone identification were performed as previously described (Nakade et al., 2014). sgRNA sequences and primers were listed in Supplemental Table S6.

#### Immunoblotting and immunofluorescence staining

IB and IF experiments were performed as described in (Xie et al., 2018); briefly, cells were washed twice with ice-cold PBS and lysed with EBC buffer (50 mM Tris (pH 7.5), 120 mM NaCl, 0.5% NP-40) with 1 mM dithiothreitol (DTT), protease inhibitors, and phosphatase inhibitors; protein concentrations were measured by Bradford or BCA reagents (Beyotime Biotechnology, China); resolved by 10 or 12% SDS-PAGE; and analyzed by immunoblotting with antibodies as indicated; For IF, cells were seeded on polylysine-coated glass coverslips and grown overnight; treated as indicated, cells were fixed with 4% paraformaldehyde (PFA) in PBS for 10 min and permeabilized with 0.1% Triton X-100 in PBS for 5 min. After blocking with 5% bovine serum albumin (BSA), cells were incubated with primary antibodies and subsequently with anti-rabbit or mouse secondary antibodies conjugated with Alexa Fluor 488 or Alexa Fluor 594. Cells were later stained with DAPI (1 µg/mL) for 5 minutes and mounted with mounting media (90% glycerol in PBS), stored at 4°C until imaging. Primary and secondary antibodies used in this study are listed in the Key resources table.

#### Live-cell microscopic imaging and quantification

Cells were grown in 4-chamber glass-bottom microwell dishes (Cellvis, D35C4-20-1-N), after transfection of mTIR plasmids as indicated, live-cell imaging was performed with a Nikon ECLIPSE Ti2 inverted microscope with a 60x oil objective. For time-lapse imaging, cells were placed in an incubation chamber maintained at 37°C with 5% CO2, exposure times were about 100 ms for mTIR, and images were taken at regular intervals. Confocal images were taken by a Zeiss LSM 880 confocal microscope with a 60x oil objective; exposure times were 50–700 ms. For cellular FRAP experiments, mTIR puncta induced by rapamycin in U2OS cells were photobleached with a 560 nm laser by Zeiss LSM880 at room temperature. The fluorescence intensities in selected regions were collected every 0.9456 s as the mean ROI, and the value of mCherry signals was normalized to the initial intensity before photobleaching. Recovery curves were plotted using GraphPad Prism software. For FRET analysis, cells were cultured in 4-chamber glass-bottom microwell dishes and transfected with mTIR plasmids as specified. Live-cell imaging was performed using a Zeiss LSM880 inverted microscope equipped with a 40x oil objective. CFP images were acquired using a 405 nm excitation wavelength and an emission range of 450 nm to 521 nm, while YFP images were acquired using a 514 nm excitation wavelength and an emission range of 540 nm to 701 nm. The traces were normalized by setting the emission ratio before drug addiction. For quantification of mTIR-positive cells, three biological replicates were photographed; about 50 cells in each sample were calculated by the percentage of mTIR-positive cells, which is defined by more than 10 visible puncta in one cell area.

#### Compound library screening and data analysis

Screening compounds were stored at 10 mM stock in DMSO. 293T cells stably expressing mTIR were seeded in 96-well glass bottom plates (P96-1.5P from Cellvis) with 8000 cells per well and grown in complete medium overnight. Plated cells were treated with compounds (10 μM), Torin1 (100 nM), or DMSO as controls in complete medium for 24 hours. Live-cell imaging was performed by Nikon ECLIPSE Ti2 inverted microscope with 60x oil objective as described above. The screening quality was evaluated by the Z’ factor using the formula:

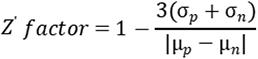

µp and σp are the mean and standard deviation values of the positive control (Torin1), µn and σn are the mean and standard deviation values of the negative control (DMSO). For the inhibitor validation experiment, the compounds were ordered separately, and the dose and treatment time were specified in each figure legend. Library compounds and other inhibitors used in this study are listed in the Key Resources Table. For quantitative analysis of screening imaging data with Image J, the puncta’s pixel fluorescence intensity and the cell’s pixel intensity were measured using the “Analyze Particle” function. The puncta signal was first masked to calculate the background cells’ pixel intensity, then the background signal was masked to calculate the puncta signals, the mTIR signal were calculated by formula :

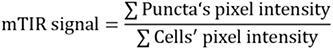

To account for variability in the data, the values obtained above were normalized based on the minimum and maximum values, which transformed the data into values ranging between 0 and 1. For further accuracy, the positive hits were manually checked and the mTIR-positive cell percentage was calculated; the negative hits were also normalized to the percentage with Torin1 as the reference, and thresholds were set at 30% to effectively eliminate the noise signals.

#### Cytosolic and nuclear protein fractionation

Fractionation was performed as described in (Rosner and Hengstschlager, 2008); briefly, cell pellets were lysed in buffer A (20 mM Tris-HCl pH 7.6, 0.6% CHAPS, 0.1 mM EDTA, and 2 mM MgCl2 supplemented with protease and phosphatase inhibitors) for 2 minutes at room temperature and for another 10 minutes on ice. Samples were then homogenized by passing through a 20-gauge needle five times. Nuclei were pelleted by centrifugation at 600 g for 5 min at 4°C, and the supernatant containing cytoplasmic proteins (Cyto) was collected and stored at -80°C. The remaining nuclei were washed three times with buffer A and lysed in buffer B (20 mM HEPES pH 7.9, 0.4 M NaCl, 2.5% glycerol, and 1 mM EDTA supplemented with protease and phosphatase inhibitors) on ice for 30 min and sonicated 3 X 10 seconds with 20% outputs. Supernatants containing soluble nucleic proteins (Nucl.) were collected by centrifugation at 20 000 g for 20 min and stored at -80°C.

#### RT-qPCR

Total RNA was extracted according to a standard RNA extraction protocol with Trizol and dissolved in DEPC-ddH2O. Reverse transcription was performed with 1 μg total RNAs using 5 X Evo M-MLV RT Reaction Mix Ver.2 kit (AG11728, Accurate Biology) according to the manufacturer’s instructions. RT-qPCR analyses were performed using the StepOnePlus with 2× RealStar Fast SYBR qPCR Mix (A303, GenStar). The quantification of mRNA was calculated with the ΔΔCt method and normalized to *β-Actin* or *UBC*. All experiments were performed in triplicate for 3 times. PCR primers were listed in Supplemental Table S6.

#### Bioinformatic analysis

The two mRNA expression cohorts were downloaded from the Gene Expression Omnibus (GEO) database under accession numbers GSE191126 and GSE64689. The DEGs between the TSA-treated and untreated groups were identified using the R package “DESeq2”. DEGs were defined using a P value less than 0.05 and an absolute log2 (fold change) larger than 0.5.

#### Quantification and statistical analysis

Image J and GraphPad Prism 9.0 were used to quantify and analyze the data and plot most of the graphics. The radar plot, bubble plot, scatter plot, and volcano plot were made by R 4.2.1 (https://www.r-project.org/). Data from biological or technical replicates are shown with a standard error of the mean (SEM). Statistical analysis was performed using a two-tailed Student’s t-test, the IC50 and *t*1/2 values were determined by fitting to a standard four-parameter logistic analysis. *P < 0.05 was considered statistically significant. All data from representative experiments were repeated at least three times independently, except those specifically noted in the figure legend.

## Supplemental items

Supplemental Figure S1. Additional characterization of mTIR, Related to Figure 1

Supplemental Figure S2. mTIR and mTIR^MT^ responded to mTORC1-mediated 4EBP1 phosphorylation, Related to Figure 2

Supplemental Figure S3. mTIR is an applicable tool for high-throughput screening and an indicator for genetic inhibition of mTORC1 signaling, Related to Figure 3

Figure S4. Stable mTIR puncta were regulated by 4EBP1 phosphatase, Related to Figure 4

Supplemental Figure S5. NmTIR responded to inhibition of PI3K/AKT/mTOR signaling in nucleus, Related to Figure 5

Supplemental Figure S6. mTIR high-throughput screening identified HDACi as mTORC1 inhibitors, Related to Figure 6

Supplemental Figure S7. HDACi inhibit mTORC1 signaling by induction of AA sensor gene expression, Related to Figure 7

Supplemental table S1. Amino acid sequences of mTIR and mutants, Related to Figure 2

Supplemental table S2. Information of Kinase inhibitors and target pathway, Related to Figure 3

Supplemental table S3. Putative 4EBP1 kinases in specific context, Related to Figure 3

Supplemental table S4. Information of positive hits from kinase screening, Related to Figure 3

Supplemental table S5. Information of positive hits from drug screening, Related to Figure 6

Supplemental table S6. DNA Oligo sequences used in this study, Related to SATR methods

Supplemental video 1. FRAP assay, Related to Figure 4

Supplemental video 2. Diffusion with AA, Related to Figure 4

Supplemental video 3. Diffusion with AA and OA, Related to Figure 4

