## Supplemental figures and table for "Living cell mTORC1 inhibition reporter mTIR reveals nutrient-sensing targets of histone deacetylase inhibitor"

SUPPLEMENTAL FIGURES & LEGENDS

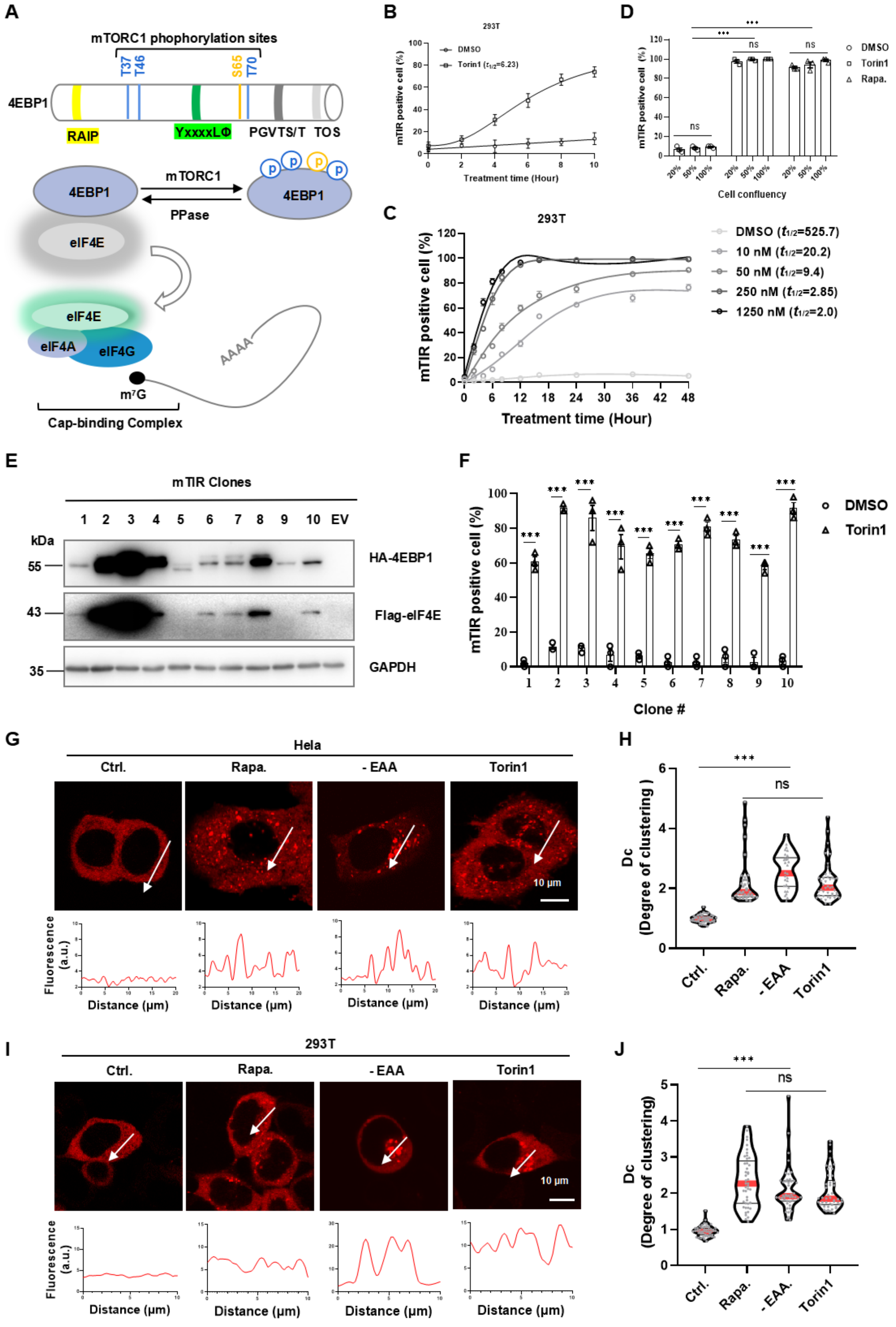


**Figure S1. Additional characterization of mTIR, Related to Figure 1**

(A) Diagram depicting the regulation of the cap-binding complex by mTORC1-mediated 4EBP1 phosphorylation in mRNA translation initiation.

(B) Time-response curve of mTIR for 100 nM Torin1 in 293T cells. $t_{1/2}$ were calculated with nonlinear fit analysis, 3 biological replicates at each time-point were calculated.

(C) Time-response curve of mTIR for different doses of Torin1 in 293T cells. $t_{1/2}$ for each dose were calculated with nonlinear fit analysis, 3 replicates at each time-point were calculated.

(D) mTIR puncta formation with different cell confluency in 293T cultures with various inhibitors.

(E) IB analysis of stable 293T clones expressing different levels of mTIR.

(F) Responses of mTIR in stable 293T clones to Torin1 inhibition (50 nM, 12 hours).

(G) (I) Representative fluorescence histograms of mTIR puncta were plotted with lines across Hela or 293T cells with indicated treatments.

(H) (J) Violin plots of Dc values in Hela and 293T cells with indicated treatments.

Data are representative of 3 independent experiments; error bars show standard error of mean (SEM) of 3 biological replicates for each treatment or time point; ns, no statistical significance, * $P < 0.05$, ** $P < 0.01$, *** $P < 0.001$, statistical analysis using two-tailed t-test; scale bar, 10 μm.

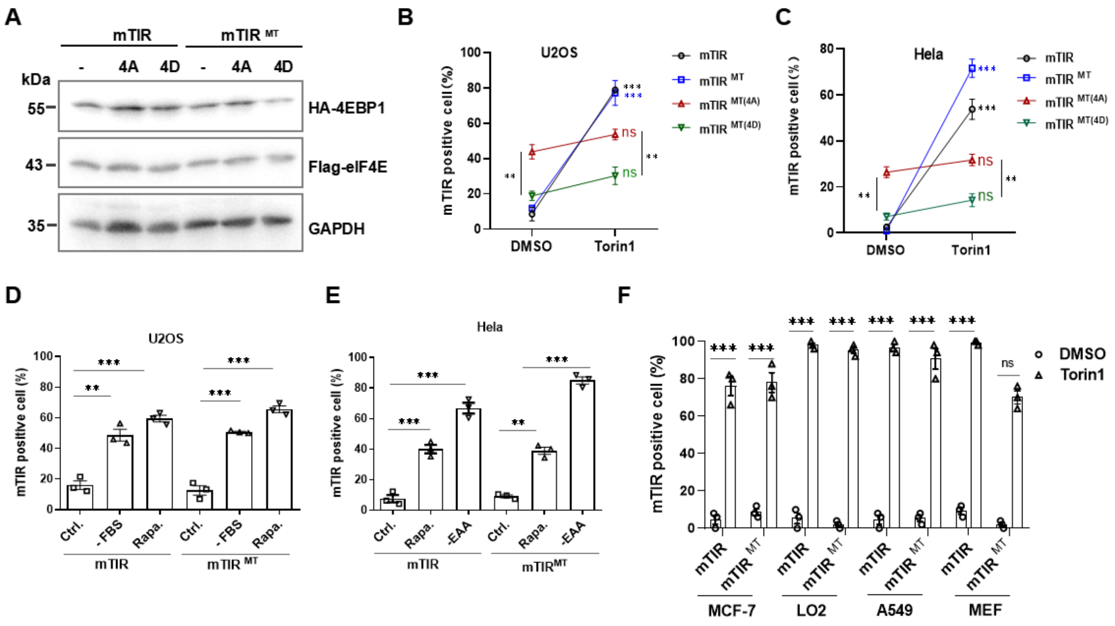


**Figure S2. mTIR and mTIR$^{MT}$ responded to mTORC1-mediated 4EBP1 phosphorylation, Related to Figure 2**

(A) IB analyzing the expression of mTIR, mTIR$^{MT}$ and its related mutants in 293T cells.

(B) (C) Responses of mTIR, mTIR$^{MT}$, and related mutants to Torin1 (50 nM, 12 hours) in U2OS and Hela cells.

(D) Responses of mTIR and mTIR$^{MT}$ to serum starvation and rapamycin treatment (100 nM, 12 hours) in U2OS cells.

(E) Responses of mTIR and mTIR$^{MT}$ to EAA starvation and rapamycin treatment (100 nM, 12 hours) in Hela cells.

(F) Responses of mTIR and mTIR$^{MT}$ to Torin1 (50 nM, 12 hours) in MEF and various human cancer cell lines.

Data are representative of 3 independent experiments; error bars show the standard error of the mean (SEM) of 3 biological replicates for each treatment or time point; ns, no statistical significance, * $P < 0.05$, ** $P < 0.01$, *** $P < 0.001$, statistical analysis using two-tailed t-test; scale bar, 10 µm.

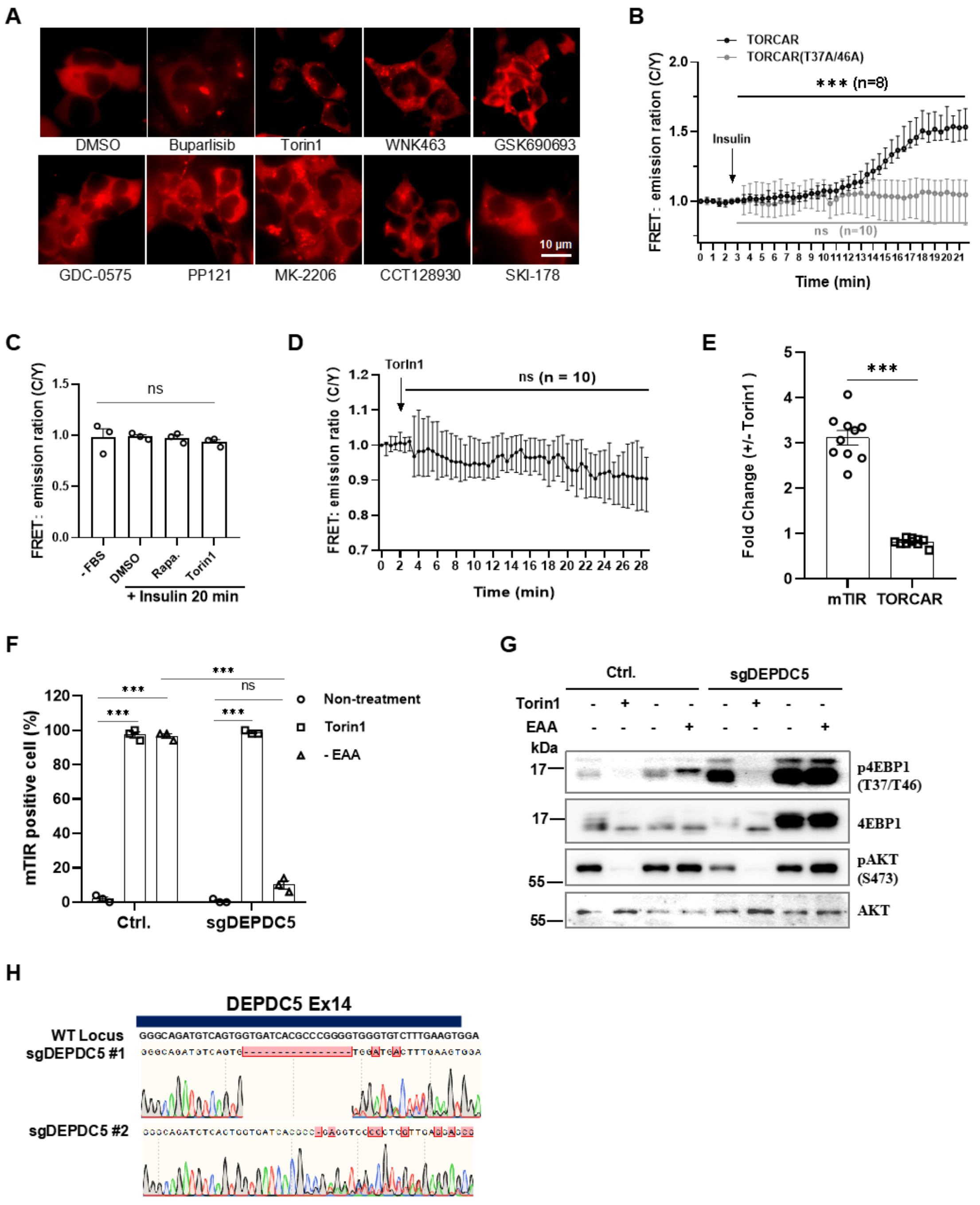


**Figure S3. mTIR is an applicable tool for high-throughput screening and an indicator for genetic inhibition of mTORC1 signaling, Related to Figure 3**

(A) Representative images of positive hits from kinase inhibitor screening.

(B) Time course of the TORCAR response to insulin (100 nM), overnight serum starved 293T cells expressing TORCAR (Black trace, n = 8) or TORCAR-T37A/46A (Grey trace, n = 10) were stimulate insulin, normalized FRET emission signal (CFP/YPET) were recorded for about 20 minutes, the FRET signals were compared between mean value of peaked level (time-point 20) and the base level (time-point 3) before insulin stimulation.

(C) TORCAR response to insulin (100 nM) with or without rapamycin and Torin1 pre-inhibition, 293T cells expressing TORCAR were treated as described in (B) except with pretreatment of rapamycin (100 nM) and Torin1(50 nM) with overnight serum starvation, the peaked FRET signals at 20 minutes were compared between different treatments.

(D) Time course of the TORCAR responses to Torin1 (50 nM) in 293T cells, FRET signals were recorded for about 30 minutes, mean value of each time point after Torin1 and the base level before Torin1 were compared.

(E) The mTIR Dc value and TORCAR FRET signal fold change comparision in 10 single cells for each reporter with Torin1 treatments in 293T cells.

(F) Quantified mTIR responses to EAA starvation with DEPDC5 depletion, wild type and DEPDC5 knockout 293T cells were treated with 50 nM Torin1 or EAA starvation for 12 hours, 3 biological replicates were photographed and quantified.

(G) IB analysis of mTORC1 signaling in response to Torin1 or EAA starvation in wild type and DEPDC5-deficient 293T cells.

(H) Identification of DEPDC5 knockout 293T clonal cells, Sanger sequencing results were shown for PCR product of DEPDC5 genomic locus in exon 14 (Ex14) covering the cut sites of Cas9/sgDEPDC5 complex.

Data are representative of 3 independent experiments; Error bars show standard error of mean (SEM) of 3 biological replicates for each treatment or time-point; ns, no statistical significance, * $P < 0.05$, ** $P < 0.01$, *** $P < 0.001$, statistical analysis using two-tailed t-test; scale bar, 10 μm.

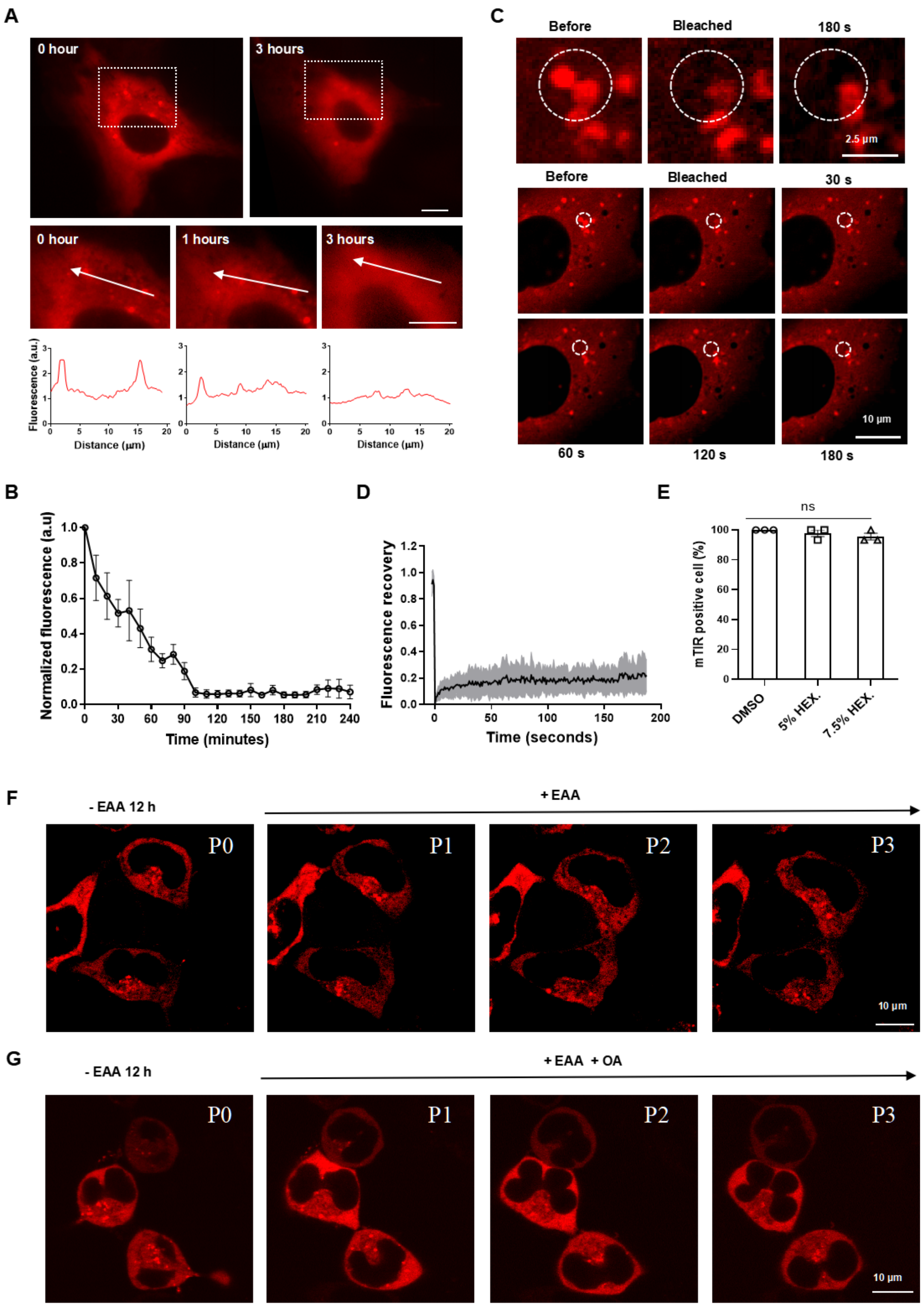


**Figure S4. Stable mTIR puncta were regulated by 4EBP1 phosphatase, Related to Figure 4**

(A) Representative time-lapse imaging of mTIR upon EAA re-stimulation in 293T cells at the indicated time point. Fluorescence histograms of mTIR puncta were plotted with lines across cells with indicated treatments; scale bar, 10 μm.

(B) Quantified fluorescence intensity for mTIR puncta upon EAA re-stimulation in 293T cells was recorded in 3 cells for 4 hours upon EAA re-stimulation, and the normalized fluorescence was calculated as the ratio of the ∑ (puncta pixel intensity) over ∑ (cells' pixel intensity) as described in the method details. Data presented as mean±SEM; experiments repeated three times.

(C) Representative FRAP images of mTIR puncta in U2OS cells. After bleaching, time-lapse images were taken at the indicated intervals for 3 minutes. The white circles showed zoomed puncta (top), and bleaching sites (bottom); scale bar, 10 µm.

(D) Quantified FRAP signals as in (C) for 10 mTIR puncta. Data plotted as mean± SEM (grey error bar, n = 10).

(E) The effect of HEX on mTIR puncta diffusion in 293T cells. mTIR puncta were induced by 50 nM Torin1 for 16 hours, then treated with the indicated HEX for 10 minutes, and photographed. mTIR-positive cells were calculated. The data are representative of three independent experiments. Error bars show the standard error of the mean (SEM) of three biological replicates for each treatment. ns, no statistical significance.

(F) (G) Representative images of mTIR over three hours upon EAA re-feeding with or without OA treatment in 293T cells. P0, P1, P2, and P3 denote 0, 1, 2, and 3 hours of EAA re-feeding, respectively. Images are representative of at least 3 independent experiments, scale bar, 10 µm.

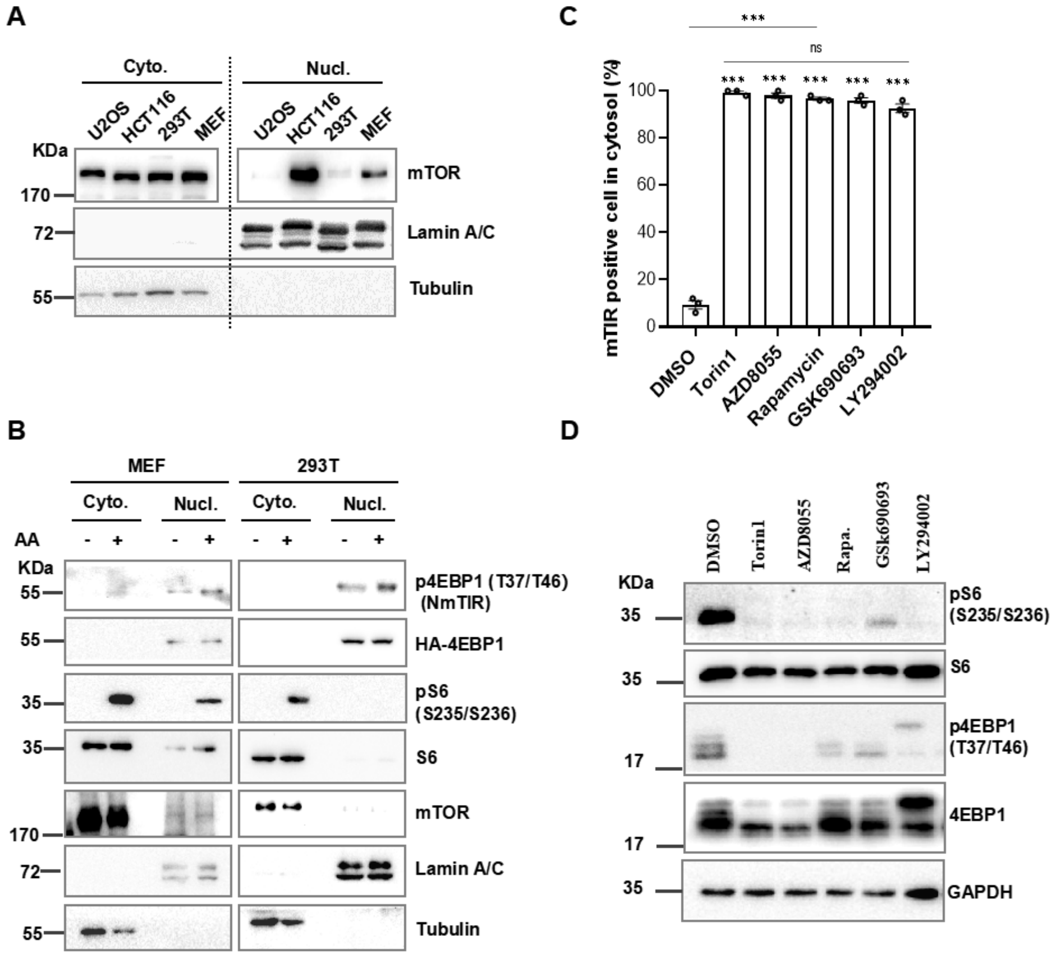


**Figure S5. NmTIR responded to inhibition of PI3K/AKT/mTOR signaling in nucleus, Related to Figure 5**

(A) IB analysis of mTOR expression in cytosolic (cyto.) and nuclear fractions (nucl.) of different cell lines.

(B) IB analysis of mTORC1 activity in the cytosolic and nuclear fractions of 293T and MEF cell lines.

(C) Responses of mTIR to inhibitors of PI3K/AKT/mTOR signaling in MEF cells as in Figure 5D. Error bars show standard error of mean (SEM) of 3 biological replicates for each treatment or time-point; *** $P < 0.001$, statistical analysis using two-tailed t-test.

(D) IB analysis of mTORC1 activity in whole cell lysate of MEF cells treated with PI3K/AKT/mTOR inhibitors as in Figure 5D.

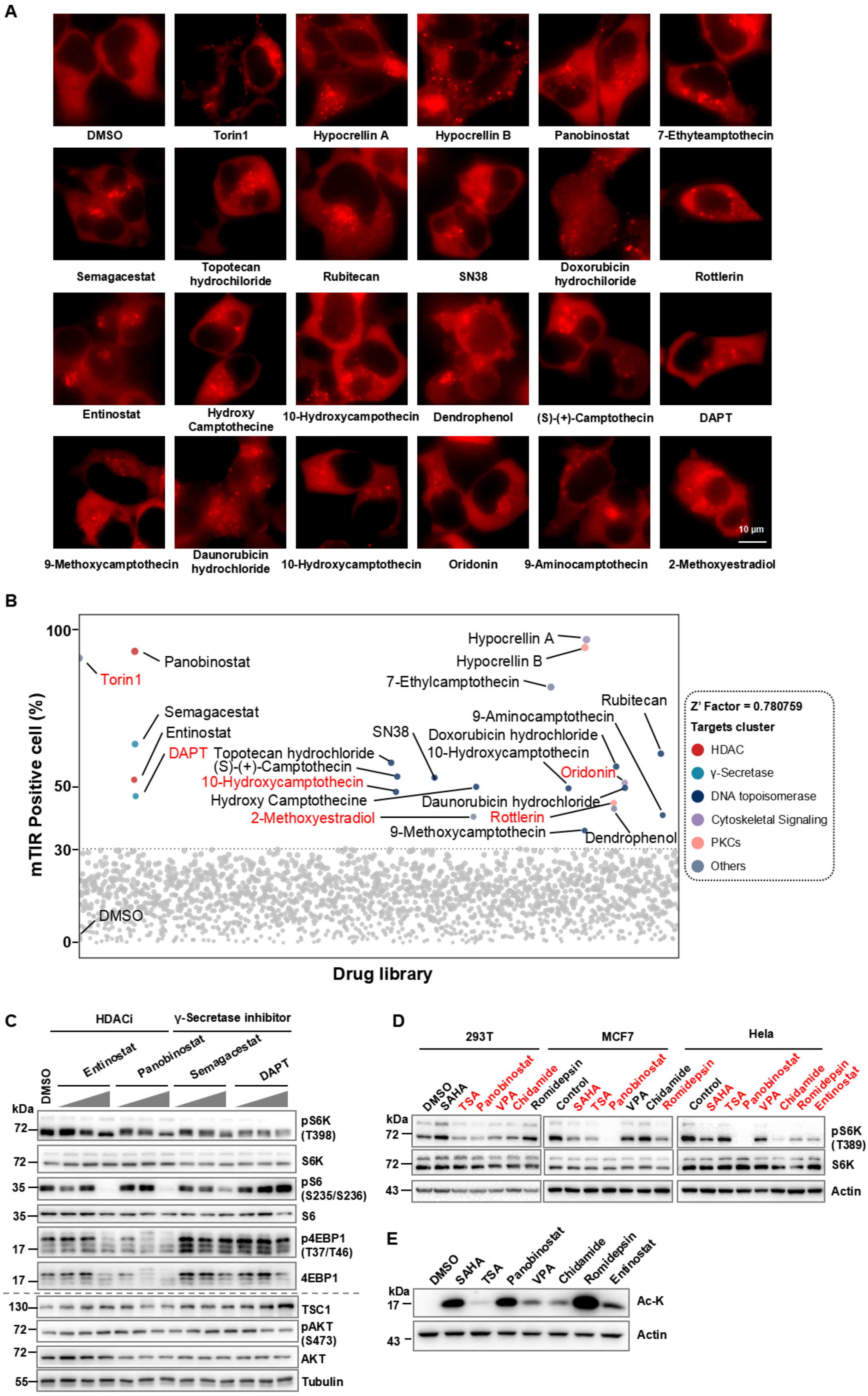
A
DMSO
Torin1
Hypocrellin A
Hypocrellin B
Panobinostat
7-Ethyteamptothecin
Semagacestat
Topotecan hydrochiloride
Rubitecan
SN38
Doxorubicin hydrochiloride
Rottlerin
Entinostat
Hydroxy Camptothecine
10-Hydroxycampothecin
Dendrophenol
(S)-(+)-Camptothecin
DAPT
9-Methoxycamptothecin
Daunorubicin hydrochiloride
10-Hydroxycamptothecin
Oridonin
9-Aminocamptothecin
2-Methoxyestradiol
10 μm
B
mTIR Positive cell (%)
100
50
30
0
Drug library
Panobinostat
Torin1
Semagacestat
Entinostat
DAPT
Topotecan hydrochloride
(S)-(+)-Camptothecin
10-Hydroxycamptothecin
Hydroxy Camptothecine
2-Methoxyestradiol
SN38
Hypocrellin A
Hypocrellin B
7-Ethylcamptothecin
Rubitecan
9-Aminocamptothecin
Doxorubicin hydrochloride
10-Hydroxycamptothecin
Oridonin
Daunorubicin hydrochloride
Rottlerin
9-Methoxycamptothecin
Dendrophenol
DMSO
Z' Factor = 0.780759
Targets cluster
HDAC
γ-Secretase
DNA topoisomerase
Cytoskeletal Signaling
PKCs
Others
C
HDACi
γ-Secretase inhibitor
DMSO
Entinostat
Panobinostat
Semagacestat
DAPT
kDa
72
35
17
130
55
pS6K (T398)
S6K
pS6 (S235/S236)
S6
p4EBP1 (T37/T46)
4EBP1
TSC1
pAKT (S473)
AKT
Tubulin
D
293T
MCF7
Hela
DMSO
Control
SAHA
TSA
Panobinostat
VPA
Chidamide
Romidepsin
Entinostat
kDa
72
43
pS6K (T389)
S6K
Actin
E
DMSO
SAHA
TSA
Panobinostat
VPA
Chidamide
Romidepsin
Entinostat
kDa
17
43
Ac-K
Actin

**Figure S6. mTIR high-throughput screening identified HDACi as mTORC1 inhibitors, Related to Figure 6**

(A) Representative images of the positive hits from 293T-mTIR live-cell screening, Hypocrellin A/B exhibit fluorescence at 680-730 nm emission overlped with mCherry, information of each hits were listed in Supplemental table S5. Scale bar, 10 µm.

(B) Scatter plots of drug library screening by mTIR reporter, details were descrbed in STAR methods, the red-highlighted drugs were found inhibiting mTORC1 in published literature, the gray dots indicate no or weak inhibitory activity while colored dots indicate identified hits, the dot line shows the 30% threshold for screening.

(C) IB analysis of mTORC1 signaling with dose-escalation (0.1, 1.0, and 10 µM for 12 hours) treatment of HDACi and γ- Secretase inhibitors in MCF7 cells.

(D) IB analysis of mTORC1 signaling with 24-hour HDACi treatment in different cell lines (10 µM SAHA, 10 µM TSA, 10 µM Panobinostat, 10 µM VPA, 10 µM Chidamide, 1.0 µM Romidepsin, and 10.0 µM Entinostat), those inhibited mTORC1 activity as indicated by S6K phosphorylation decrease were red highlighted.

(E) IB analysis of acetylated histones in response to HDACi in 293T cells as described in (D).

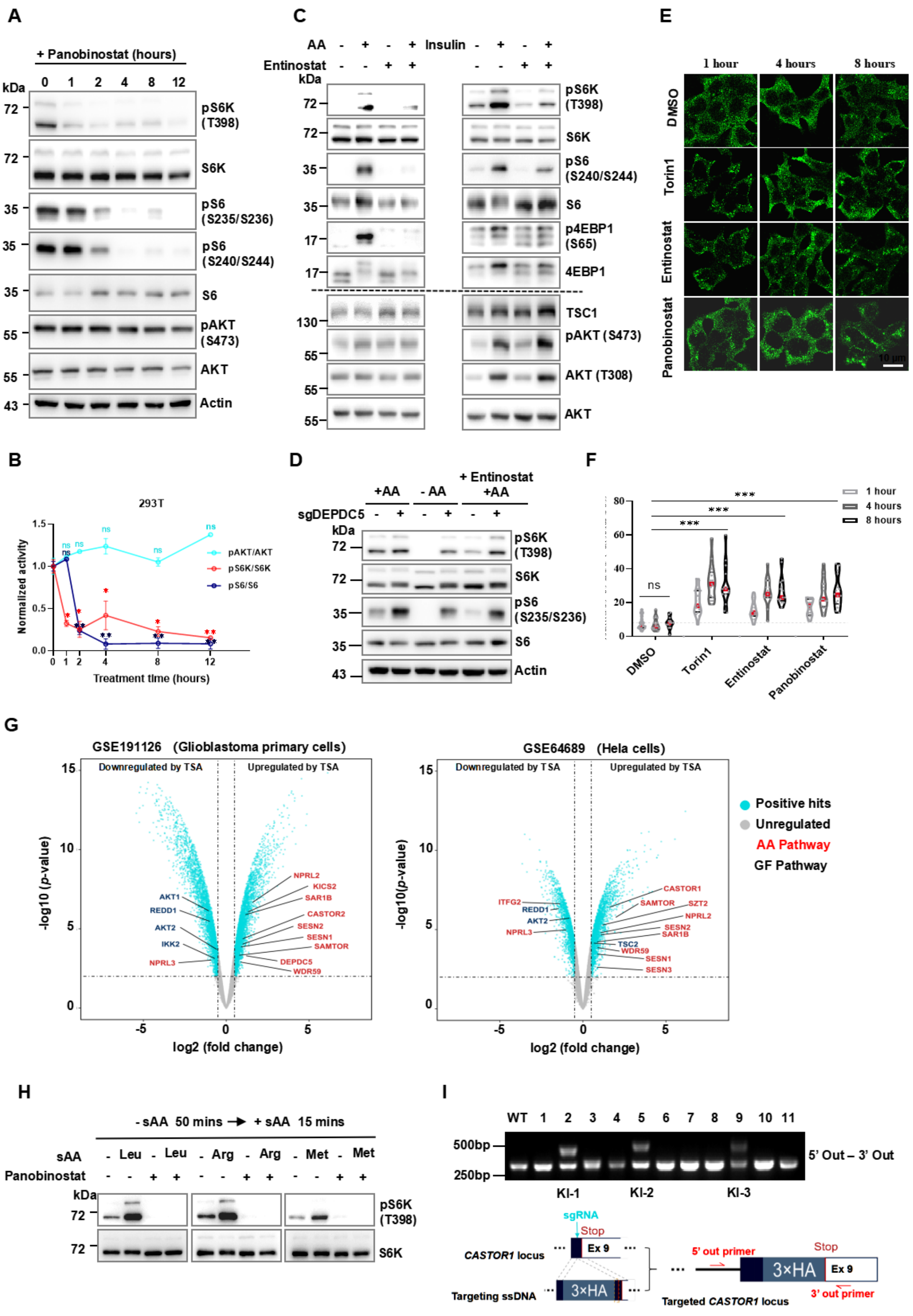
A
+ Panobinostat (hours)
kDa
0 1 2 4 8 12
72
pS6K (T398)
S6K
35
pS6 (S235/S236)
pS6 (S240/S244)
S6
55
pAKT (S473)
AKT
43
Actin
C
AA - + - +
Entinostat - - + +
Insulin - + - +
kDa
72
35
17
130
55
pS6K (T398)
S6K
pS6 (S240/S244)
S6
p4EBP1 (S65)
4EBP1
TSC1
pAKT (S473)
AKT (T308)
AKT
E
1 hour
4 hours
8 hours
DMSO
Torin1
Entinostat
Panobinostat
10 µm
B
293T
pAKT/AKT
pS6K/S6K
pS6/S6
Normalized activity
Treatment time (hours)
ns
D
+AA
- AA
+ Entinostat +AA
sgDEPDC5
kDa
72
35
43
pS6K (T398)
S6K
pS6 (S235/S236)
S6
Actin
F
1 hour
4 hours
8 hours
ns
***
DMSO
Torin1
Entinostat
Panobinostat
G
GSE191126 (Glioblastoma primary cells)
Downregulated by TSA
Upregulated by TSA
-log10 (p-value)
log2 (fold change)
AKT1
REDD1
AKT2
IKK2
NPRL3
NPRL2
KICS2
SAR1B
CASTOR2
SESN2
SESN1
SAMTOR
DEPDC5
WDR59
GSE64689 (Hela cells)
ITFG2
CASTOR1
SZT2
TSC2
SESN3
Positive hits
Unregulated
AA Pathway
GF Pathway
H
- sAA 50 mins → + sAA 15 mins
sAA
Leu
Arg
Met
Panobinostat
pS6K (T398)
S6K
I
WT 1 2 3 4 5 6 7 8 9 10 11
500bp
250bp
5' Out – 3' Out
KI-1
KI-2
KI-3
sgRNA
Stop
Ex 9
CASTOR1 locus
Targeting ssDNA
3×HA
5' out primer
3' out primer
Targeted CASTOR1 locus

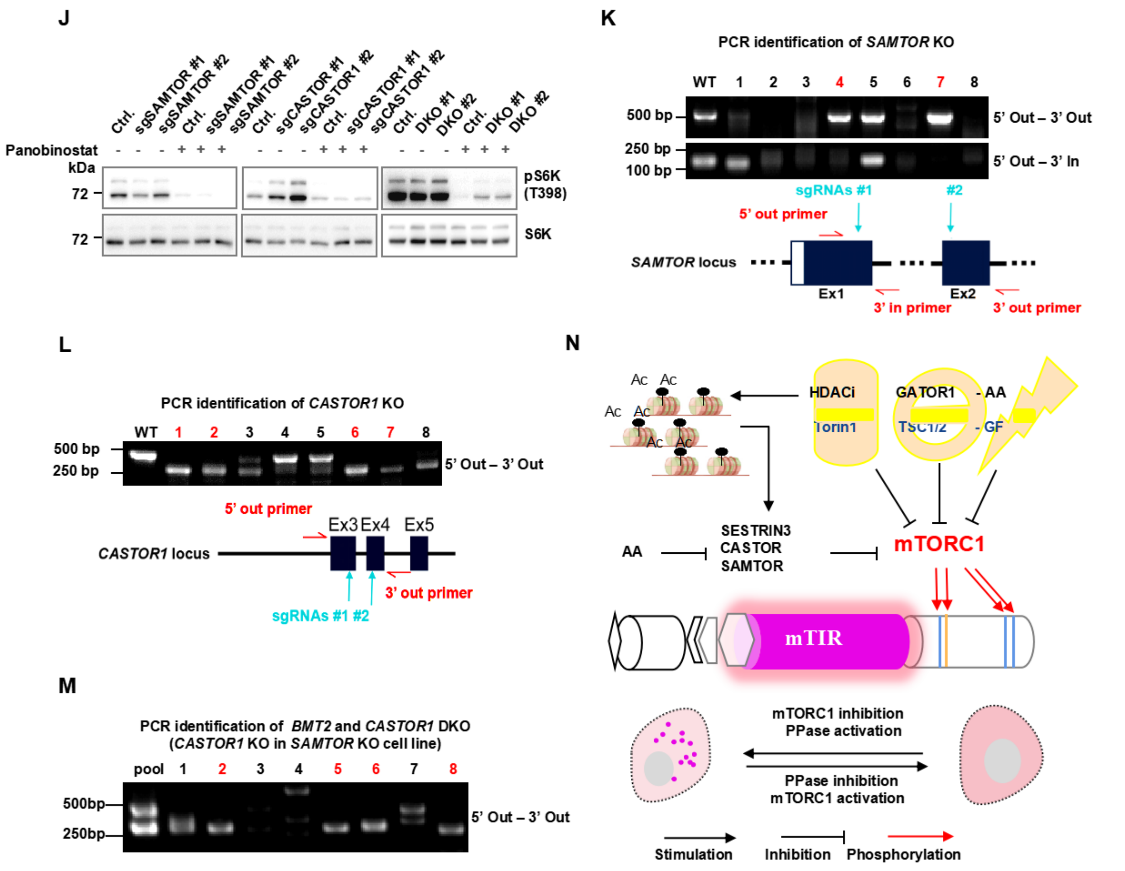


**Figure S7. HDACi inhibit mTORC1 signaling by induction of AA sensor gene expression, Related to Figure 7**

(A) IB analysis of mTOR activity in response to time course Panobinostat (10 μM) inhibition in 293T cells.

(B) Normalized mTOR activity (pS6K, pS6 and pAKT were normalized to total S6K, S6 and AKT protein level) with time course Panobinostat inhibition in 2 independent experiments.

(C) IB analysis of mTOR activity in response to Entinostat with AA or inuslin stimulation, 293T cells were pre-treated with DMSO or 10 μM Entinostat for 12 hours, then starved with AA for 50 minutes (min) and stimulated with AA for 15 min, or serum starved for 12 hours and stimulated with 100 nM insulin for 15 min.

(D) IB analysis of mTORC1 activity with 10 μM Entinostat inhibition in DEPDC5-depleted 293T cells with or without AA stimulation.

(E) Time-course autophagy analysis by IF staining with the LC3 A/B antibody in response to HDACi and Torin1 in 293T cells treated with DMSO, Torin1 (50 nM), Entinostat (10 μM), or Panobinostat (10 μM). Scale bar, 10 μm.

(F) Violin plots of quantified LC3 A/B puncta per cell in (E). Red line, median; black line, interquartile ranges, 15-20 cells were quantified from each of three biological replicates.

(G) Volcano plot depicting differentially expressed genes from TSA treated and un-treated sample. colored dots represent positive hits defined by P value less than 0.05 and log2 fold change larger than 0.5, labeled points denote mTOR-related genes in amino acid pathway (Red) and grow factor pathway (Blue).

(H) IB analysis of Panobinostat inhibition (10 μM, 12hours) of mTORC1 activity in 293T cells with 2 hours of single AA starvation (sAA) and following 15 minutes of sAA stimulation: Leu (400 μM), Arg (600 μM) and Met (100 μM).
(I) 293T clonal identification strategy for 3xHA endogenously tagged CASTOR1 gene loci, properly targeted 3xHA sequence was identified by out-out primers flanked in the last exon, the upper band PCR product showed 3xHA tagged locus, and the lower band showed the wild type locus.
(J) IB analysis of mTORC1 inhibition by Panobinostat (10 μM, 12hours) in SAMTOR, CASTOR1 or double knockout (DKO) 293T cells under normal culture conditions.
(K) (L) PCR identification strategy for SAMTOR and CASTOR1 gene knockout in 293T clones with two sgRNA.
(M) Schematic diagram of mTIR reporter and its regulation by inhibitors, GAP proteins, and physiological stresses, HDACi-induced AA sensor gene expression (SESTRIN3, CASTOR, and SAMTOR) specifically inhibit mTORC1, which together with protein phosphatase (PPase) leads to 4EBP1 dephosphorylation and mTIR puncta formation, this process could be reversed by PPase inhibition and mTORC1 re-activation.

**Supplemental table S1. Amino acid sequences of mTIR and mutants, Related to Figure 2**

| | HA Tag-4EBP1-mCherry-Linker-HOTag3-P2A-Flag Tag-eIF4E-Linker-HOTag6 |
|---|---|
| **mTIR** | MYPYDVPDYAMSGGSSCSQTPSRAIPATRRVVLGDGVQLPPGDYSTTPGGTLFSTTPG<br>GTRIIYDRKFLMECRNSPVTKTPPRDLPTIPGVTSPSSDEPPMEASQSHLRNSPEDKRAG<br>GEESQFEMDIGGSGSGGGTPVATMVSKGEEDNMAIIKEFMRFKVHMEGSVNGHEFEIE<br>GEGEGRPYEGTQTAKLKVTKGGPLPFAWDILSPQFMYGSKAYVKHPADIPDYLKLSFPE<br>GFKWERVMNFEDGGVVTVTQDSSLQDGEFIYKVKLRGTNFPSDGPVMQKKTMGWEA<br>SSERMYPEDGALKGEIKQRLKLKDGGHYDAEVKTTYKAKKPVQLPGAYNVNIKLDITSH<br>NEDYTIVEQYERAEGRHSTGGMDELYKSGLRSGSGSAGGSAGGSAGGSAGGSAGGSAG<br>GSAGGSRGEIAKSLKEIAKSLKEIAWSLKEIAKSLKGGTEGRGSLLTCGDVEENPGPKLDYK<br>DDDDKMATVEPETTPTPNPPTTEEEKTESNQEVANPEHYIKHPLQNRWALWFFKNDKS<br>KTWQANLRLISKFDTVEDFWALYNHIQLSSNLMPGCDYSLFKDGIEPMWEDEKNKRGG<br>RWLITLNKQQRRSDLDRFWLETLLCLIGESFDDYSDDVCGAVVNVRAKGDKIAIWTTEC<br>ENREAVTHIGRVYKERLGLPPKIVIGYQSHADTATKSGSTTKNRFVVVDGSGSAGGSAGG<br>SAGGSAGGSAGGSAGGSAGGSRTLREIEELLRKIIEDSVRSVAELEDIEKWLKKI* |
| **mTIR$^{(4A)}$** | MYPYDVPDYAMSGGSSCSQTPSRAIPATRRVVLGDGVQLPPGDYSTAPGGTLFSTAPG<br>GTRIIYDRKFLMECRNAPVTKAPPRDLPTIPGVTSPSSDEPPMEASQSHLRNSPEDKRAG<br>GEESQFEMDIGGSGSGGGTPVATMVSKGEEDNMAIIKEFMRFKVHMEGSVNGHEFEIE<br>GEGEGRPYEGTQTAKLKVTKGGPLPFAWDILSPQFMYGSKAYVKHPADIPDYLKLSFPE<br>GFKWERVMNFEDGGVVTVTQDSSLQDGEFIYKVKLRGTNFPSDGPVMQKKTMGWEA<br>SSERMYPEDGALKGEIKQRLKLKDGGHYDAEVKTTYKAKKPVQLPGAYNVNIKLDITSH<br>NEDYTIVEQYERAEGRHSTGGMDELYKSGLRSGSGSAGGSAGGSAGGSAGGSAGGSAG<br>GSAGGSRGEIAKSLKEIAKSLKEIAWSLKEIAKSLKGGTEGRGSLLTCGDVEENPGPKLDYK<br>DDDDKMATVEPETTPTPNPPTTEEEKTESNQEVANPEHYIKHPLQNRWALWFFKNDKS<br>KTWQANLRLISKFDTVEDFWALYNHIQLSSNLMPGCDYSLFKDGIEPMWEDEKNKRGG<br>RWLITLNKQQRRSDLDRFWLETLLCLIGESFDDYSDDVCGAVVNVRAKGDKIAIWTTEC<br>ENREAVTHIGRVYKERLGLPPKIVIGYQSHADTATKSGSTTKNRFVVVDGSGSAGGSAGG<br>SAGGSAGGSAGGSAGGSAGGSRTLREIEELLRKIIEDSVRSVAELEDIEKWLKKI* |
| **mTIR$^{(4D)}$** | MYPYDVPDYAMSGGSSCSQTPSRAIPATRRVVLGDGVQLPPGDYSTDPGGTLFSTDPG<br>GTRIIYDRKFLMECRNDPVTKDPPRDLPTIPGVTSPSSDEPPMEASQSHLRNSPEDKRAG<br>GEESQFEMDIGGSGSGGGTPVATMVSKGEEDNMAIIKEFMRFKVHMEGSVNGHEFEIE<br>GEGEGRPYEGTQTAKLKVTKGGPLPFAWDILSPQFMYGSKAYVKHPADIPDYLKLSFPE<br>GFKWERVMNFEDGGVVTVTQDSSLQDGEFIYKVKLRGTNFPSDGPVMQKKTMGWEA<br>SSERMYPEDGALKGEIKQRLKLKDGGHYDAEVKTTYKAKKPVQLPGAYNVNIKLDITSH<br>NEDYTIVEQYERAEGRHSTGGMDELYKSGLRSGSGSAGGSAGGSAGGSAGGSAGGSAG<br>GSAGGSRGEIAKSLKEIAKSLKEIAWSLKEIAKSLKGGTEGRGSLLTCGDVEENPGPKLDYK<br>DDDDKMATVEPETTPTPNPPTTEEEKTESNQEVANPEHYIKHPLQNRWALWFFKNDKS<br>KTWQANLRLISKFDTVEDFWALYNHIQLSSNLMPGCDYSLFKDGIEPMWEDEKNKRGG<br>RWLITLNKQQRRSDLDRFWLETLLCLIGESFDDYSDDVCGAVVNVRAKGDKIAIWTTEC<br>ENREAVTHIGRVYKERLGLPPKIVIGYQSHADTATKSGSTTKNRFVVVDGSGSAGGSAGG<br>SAGGSAGGSAGGSAGGSAGGSRTLREIEELLRKIIEDSVRSVAELEDIEKWLKKI* |

| | |
|---|---|
| **mTIR$^{MT}$** | **MYPYDVPDYAMAGGAACAQAPARAIPAARRVVLGDGVQLPPGDYAATPGGALFAATPGGARIIYDRKFLMECRNSPVAKTPPRDLPAIPGVAAPAADEPPMEAAQAHLRNAPEDKRAGGEEAQFEMDI**GGSGSGGGTPVAT**MVSKGEEDNMAIIKEFMRFKVHMEGSVNGHEFEIEGEGEGRPYEGTQTAKLKVTKGGPLPFAWDILSPQFMYGSKAYVKHPADIPDYLKLSFPEGFKWERVMNFEDGGVVTVTQDSSLQDGEFIYKVKLRGTNFPSDGPVMQKKTMGWEASSERMYPEDGALKGEIKQRLKLKDGGHYDAEVKTTYKAKKPVQLPGAYNVNIKLDITSHNEDYTIVEQYERAEGRHSTGGMDELYK**SGLRSGSGSAGGSAGGSAGGSAGGSAGGSAGGSAGGSR**GEIAKSLKEIAKSLKEIAWSLKEIAKSLKGGTEGRGSLLTCGDVEENPGP**KL**DYKDDDDKMATVEPETTPTPNPPTTEEEKTESNQEVANPEHYIKHPLQNRWALWFFKNDKSKTWQANLRLISKFDTVEDFWALYNHIQLSSNLMPGCDYSLFKDGIEPMWEDEKNKRGGRWLITLNKQQRRSDLDRFWLETLLCLIGESFDDYSDDVCGAVVNVRAKGDKIAIWTTECENREAVTHIGRVYKERLGLPPKIVIGYQSHADTATKSGSTTKNRFVV**VDGSGSAGGSAGGSAGGSAGGSAGGSAGGSAGGSR**TLREIEELLRKIIEDSVRSVAELEDIEKWLKKI*** |
| **mTIR$^{MT(4A)}$** | **MYPYDVPDYAMAGGAACAQAPARAIPAARRVVLGDGVQLPPGDYAAAPGGALFAAAPGGARIIYDRKFLMECRNAPVAKAPPRDLPAIPGVAAPAADEPPMEAAQAHLRNAPEDKRAGGEEAQFEMDI**GGSGSGGGTPVAT**MVSKGEEDNMAIIKEFMRFKVHMEGSVNGHEFEIEGEGEGRPYEGTQTAKLKVTKGGPLPFAWDILSPQFMYGSKAYVKHPADIPDYLKLSFPEGFKWERVMNFEDGGVVTVTQDSSLQDGEFIYKVKLRGTNFPSDGPVMQKKTMGWEASSERMYPEDGALKGEIKQRLKLKDGGHYDAEVKTTYKAKKPVQLPGAYNVNIKLDITSHNEDYTIVEQYERAEGRHSTGGMDELYK**SGLRSGSGSAGGSAGGSAGGSAGGSAGGSAGGSAGGSR**GEIAKSLKEIAKSLKEIAWSLKEIAKSLKGGTEGRGSLLTCGDVEENPGP**KL**DYKDDDDKMATVEPETTPTPNPPTTEEEKTESNQEVANPEHYIKHPLQNRWALWFFKNDKSKTWQANLRLISKFDTVEDFWALYNHIQLSSNLMPGCDYSLFKDGIEPMWEDEKNKRGGRWLITLNKQQRRSDLDRFWLETLLCLIGESFDDYSDDVCGAVVNVRAKGDKIAIWTTECENREAVTHIGRVYKERLGLPPKIVIGYQSHADTATKSGSTTKNRFVV**VDGSGSAGGSAGGSAGGSAGGSAGGSAGGSAGGSR**TLREIEELLRKIIEDSVRSVAELEDIEKWLKKI*** |
| **mTIR$^{MT(4D)}$** | **MYPYDVPDYAMAGGAACAQAPARAIPAARRVVLGDGVQLPPGDYAADPGGALFAADPGGARIIYDRKFLMECRNDPVAKDPPRDLPAIPGVAAPAADEPPMEAAQAHLRNAPEDKRAGGEEAQFEMDI**GGSGSGGGTPVAT**MVSKGEEDNMAIIKEFMRFKVHMEGSVNGHEFEIEGEGEGRPYEGTQTAKLKVTKGGPLPFAWDILSPQFMYGSKAYVKHPADIPDYLKLSFPEGFKWERVMNFEDGGVVTVTQDSSLQDGEFIYKVKLRGTNFPSDGPVMQKKTMGWEASSERMYPEDGALKGEIKQRLKLKDGGHYDAEVKTTYKAKKPVQLPGAYNVNIKLDITSHNEDYTIVEQYERAEGRHSTGGMDELYK**SGLRSGSGSAGGSAGGSAGGSAGGSAGGSAGGSAGGSR**GEIAKSLKEIAKSLKEIAWSLKEIAKSLKGGTEGRGSLLTCGDVEENPGP**KL**DYKDDDDKMATVEPETTPTPNPPTTEEEKTESNQEVANPEHYIKHPLQNRWALWFFKNDKSKTWQANLRLISKFDTVEDFWALYNHIQLSSNLMPGCDYSLFKDGIEPMWEDEKNKRGGRWLITLNKQQRRSDLDRFWLETLLCLIGESFDDYSDDVCGAVVNVRAKGDKIAIWTTECENREAVTHIGRVYKERLGLPPKIVIGYQSHADTATKSGSTTKNRFVV**VDGSGSAGGSAGGSAGGSAGGSAGGSAGGSAGGSR**TLREIEELLRKIIEDSVRSVAELEDIEKWLKKI*** |

**Supplemental table S2. Information of Kinase inhibitors and target pathway, Related to Figure 3**

| target pathway | Number of inhibitors |
|---|---|
| Cell Cycle | 59 |
| GPCR/G Protein | 4 |
| JAK/STAT Signaling | 7 |
| MAPK/ERK Pathway | 28 |
| Membrane Transporter/Ion Channel | 3 |
| NF-κB | 4 |
| PI3K/Akt/mTOR | 9 |
| Protein Tyrosine Kinase/RTK | 61 |
| Stem Cell/Wnt | 20 |
| TGF-β/Smad | 8 |
| Other unclassified | 48 |

**Supplemental table S3. Putative 4EBP1 kinases in specific context , Related to Figure 3**

| Kinase name | Phosphorylation-sites | Dependence on mTOR | Phosphorylation context | Reference |
|---|---|---|---|---|
| AKT | ND. | Yes | GF stimulation | (Anne-Claude Gingras, 1997; Gingras et al., 1998) |
| ATM | S111 | ND | GF stimulation | (Yang and Kastan, 2000) |
| Pim2 | T37/T46/S65/T70 | ND | GF stimulation | (Fox et al., 2003) |
| ERK1/2 | S65/T70 | Yes | TPA stimulated | (Herbert et al., 2002) |
| CDK1 | T37/T46/S65/S83/S101 | ND | Cell cycle-dependant | (Celestino Velásquez, 2016; Greenberg and Zimmer, 2005; Shuda et al., 2015; Velasquez et al., 2016) |
| CDK4 | T37/T46/S65/S101 | ND | Cell cycle-dependant | (Mitchell et al., 2019, 2020) |
| CDK12 | S65/T70 | Yes | Cell cycle-dependant | (Choi et al., 2019) |
| PLK1 | T37/T46 | ND | Paclitaxel induced cell cycle arrest | (Severance and Latham, 2017; Shang et al., 2012) |
| p38/MSK | T37/T46/S65/T70 | ND | UVB induced DNA Damage | (Liu et al., 2002) |
| LRRK2 | T37/T46 | ND | Oxidative stress | (Imai et al., 2008) |
| CK1ε | T41/T50 | ND | Specified in breast cancer cell lines | (Shin et al., 2014a) |
| GSK3β | T37/T46/S6/Thr 70 | ND | Specified in some cancer cell lines | (Shin et al., 2014b) |

**Supplemental table S4. Information of positive hits from kinase screening, Related to Figure 3**

| Inhibitor | Target | Relation to mTOR | Reference | Puncta cell (%) |
|---|---|---|---|---|
| Buparlisib | PI3K | PI3K/AKT/mTOR pathway | (Burger et al., 2011; McPherson et al., 2020) | 97.06197 |
| Torin1 | mTOR | PI3K/AKT/mTOR pathway | (Thoreen et al., 2009) | 85.3 |
| GSK690693 | AKT/AMPK | PI3K/AKT/mTOR pathway | (Altomare et al., 2010; Rhodes et al., 2008) | 60 |
| PP121 | mTOR/PDGFR | PI3K/AKT/mTOR pathway | (Apsel et al., 2008) | 49.84095 |
| MK-2206 | AKT | PI3K/AKT/mTOR pathway | (Hirai et al., 2010) | 39.68765 |
| CCT128930 | AKT | PI3K/AKT/mTOR pathway | (Yap et al., 2011) | 38.37867 |
| GDC-0575 | CHK1 | Genotoxic stress inhibit mTORC1 | (Ma et al., 2018) | 55 |
| WNK463 | WNKs | mTOR regulating kinase | (Liu et al., 2022) | 64.46429 |
| SKI-178 | SPHK | mTOR regulating kinase | (Jesko et al., 2019; Kim et al., 2018) | 38.33333 |

**Supplemental table S5. Information of positive hits from drug screening, Related to Figure 6**

| Drug | Target | Relation to mTOR | Reference | Puncta cell (%) |
|---|---|---|---|---|
| Entinostat | HDAC | ND | ND | 52.08333 |
| Panobinostat | HDAC | ND | ND | 93.09524 |
| Semagacestat | Y-secretase | ND | ND | 63.47882 |
| DAPT | Y-Secretase | mTORC1 inhibition | (Song et al., 2015) | 46.82821 |
| 2-Methoxyestradiol | microtubule/HIF-1α/2α | mTORC1 inhibition | (Zhou et al., 2018) | 40.24123 |
| Rottlerin | PKC | mTORC1 inhibition | (Balgi et al., 2009; Daveri et al., 2016; Torricelli et al., 2015) | 44.67019 |
| Oridonin | AKT | mTORC1 inhibition | (Wang et al., 2014) | 51.07415 |
| Dendrophenol | NF-κB | ND | ND | 42.80423 |
| Hypocrellin B | Unknown | Autofluorescence/ND | (Xu et al., 2004) | 94.3105 |
| Hypocrellin A | PKC/others | Autofluorescence/ND | (Xu et al., 2004) | 96.82111 |
| Topotecan hydrochloride | Topoisomerase I | mTORC1 inhibition | (Ma et al., 2018) | 57.51355 |
| 10-Hydroxycamptothecin | Topoisomerase I | mTORC1 inhibition | (Ma et al., 2018) | 48.18713 |
| (S)-(+)-Camptothecin | Topoisomerase I | mTORC1 inhibition | (Ma et al., 2018) | 53.09106 |
| SN38 | Topoisomerase I, | mTORC1 inhibition | (Ma et al., 2018) | 52.73268 |
| Hydroxy Camptothecine | Topoisomerase I | mTORC1 inhibition | (Ma et al., 2018) | 49.80495 |
| 7-Ethylcamptothecin | Topoisomerase I | mTORC1 inhibition | (Ma et al., 2018) | 81.66667 |
| 10-Hydroxycamptothecin | Topoisomerase I | mTORC1 inhibition | (Ma et al., 2018) | 49.32921 |
| 9-Methoxycamptothecin | Topoisomerase I | mTORC1 inhibition | (Ma et al., 2018) | 35.83057 |

| | | | | |
|---|---|---|---|---|
| Doxorubicin hydrochloride | Topoisomerase I | mTORC1 inhibition | (Ma et al., 2018) | 56.28816 |
| Daunorubicin hydrochloride | Topoisomerase II | mTORC1 inhibition | (Ma et al., 2018) | 49.45773 |
| Rubitecan | Topoisomerase I | mTORC1 inhibition | (Ma et al., 2018) | 60.401 |
| 9-Aminocamptothecin | Topoisomerase I | mTORC1 inhibition | (Ma et al., 2018) | 40.76479 |

**Supplemental table S6. DNA Oligo sequences used in this study, Related to SATR methods**

| **Primers used in quantitative RT-PCR** | | |
|---|---|---|
| **Gene** | **Primer** | **Sequence (5'-3')** |
| *NPRL2* | Forward | AAAGAAGCTGATCGGCTGTCC |
| | Reverse | GCTCTCTAGCTCTAGTGTGGT |
| *DEPDC5* | Forward | AGTGTTCCGGCTGAGACCTTA |
| | Reverse | CCACGGCCAATATACTGATCCT |
| *SESN1* | Forward | TGCTTTGGGCCGTTTGGATAA |
| | Reverse | TGTAGTGACGATAATGTAGGGGT |
| *SESN2* | Forward | TCTTACCTGGTAGGCTCCCAC |
| | Reverse | AGCAACTTGTTGATCTCGCTG |
| *SESN3* | Forward | ACCTGCTCTGTACCAACTGC |
| | Reverse | GACGACCGGATGTAGAGTATTCT |
| *SAR1B* | Forward | TACAGTGGTTTCAGCAGTGTG |
| | Reverse | AGTGGGATGTAATGTTGGGACA |
| *CASTOR1* | Forward | TCGCCACCACCCTCATAGAT |
| | Reverse | AGGTCACTGGGGAACTTTTTCT |
| *CASTOR2* | Forward | AGGAAGGATTCCTAGAGCTGC |
| | Reverse | AGTGGGGCGATGACTGACTT |
| *SAMTOR* | Forward | AGAAGTACCGAGAAGTGGGAG |
| | Reverse | ACCTTCGCCCTCACAAGTTT |
| *PTEN* | Forward | TTTGAAGACCATAACCCACCAC |
| | Reverse | ATTACACCAGTTCGTCCCTTTC |
| *TSC1* | Forward | CAACAAGCAAATGTCGGGGAG |
| | Reverse | CATAGGGCCACGGTCAGAA |
| *TSC2* | Forward | ATAGCTGTTACCTCGACGAGT |
| | Reverse | TGCAGGGAGACCTCTATGTCC |
| *REDD1* | Forward | TGGGCAAAGAACTACTGCG |
| | Reverse | AGAGTTGGCGGAGCTAAACAG |
| *ACTB* | Forward | CATGTACGTTGCTATCCAGGC |
| | Reverse | CTCCTTAATGTCACGCACGAT |
| *UBC* | Forward | ATTTGGGTCGCGGTTCTTG |
| | Reverse | TGCCTTGACATTCTCGATGGT |
| **sgRNA sequences** | | |
| **Gene** | **Primer** | **Sequence (5'-3')** |
| sgDEPDC5#1 | Forward | CACCGGTCAGTGGTGATCACGCCCG |
| | Reverse | AAACCGGGCGTGATCACCACTGACC |
| sgDEPDC5#2 | Forward | CACCGTGTTAATGTCGTAGACCCTA |
| | Reverse | AAACTAGGGTCTACGACATTAACA |
| sgCASTOR1#1 | Forward | ACGCGTCTCACACCGGACACGTGGTGCTCGGCCAGGTTTTAGAGCTAG |

| | | |
|---|---|---|
| | | AAATAGCAAG |
| | Reverse | ACGCGTCTCAAAACGCGTGTGGATCACCACGGACCGGTGTTTCGTCCTTTCCAC |
| sgSAMTOR#1 | Forward | ACGCGTCTCACACCGGGTACTTCTTGCGGAGCCGCGTTTTAGAGCTAGAAATAGCAAG |
| | Reverse | ACGCGTCTCAAAACCACAGTGTTCTCGCCAGATCCGGTGTTTCGTCCTTTCCAC |
| sgCASTOR1#2 (KI) | Forward | CACCGGTCCTCCAGCGGCGGCAGGA |
| | Reverse | AAACTCCTGCCGCCGCTGGAGGACC |
| **Primers used in point mutation and subcloning** | | |
| **Gene** | **Primer** | **Sequences (5'-3')** |
| *4EBP1* | Forward | GGTTAAGCTTGACTACAAGGACGATGACGA |
| | Reverse | GGTTACCGGTGTGCCTCCACCGCTGCCAGATCCGCCAATGTCCATCTCAAACTGTG |
| *4EBP1*(S65A/T70A) | Forward | ATGGAGTGTCGGAACGCCCCTGTGACCAAAGCCCCCCCAAGGGATCTG |
| | Reverse | CAGATCCCTTGGGGGGGCTTTGGTCACAGGGGCGTTCCGACACTCCAT |
| *4EBP1*(S65D/T70D) | Forward | CTGATGGAGTGTCGGAACGACCCTGTGACCAAAGACCCCCCAAGGGATCTG |
| | Reverse | CAGATCCCTTGGGGGGTCTTTGGTCACAGGGTCGTTCCGACACTCCATCAG |
| *4EBP1*(T37A/T46A) | Forward | GGGGACTACAGCACGGCCCCCGGCGGCACGCTCTTCAGCACCGCCCCGGGAGGTACCAGG |
| | Reverse | CCTGGTACCTCCCGGGGCGGTGCTGAAGAGCGTGCCGCCGGGGGCCGTGCTGTAGTCCCC |
| *4EBP1*(T37D/T46D) | Forward | GGGGACTACAGCACGGACCCCGGCGGCACGCTCTTCAGCACCGACCCGGGAGGTACCAGG |
| | Reverse | CCTGGTACCTCCCGGGTCGGTGCTGAAGAGCGTGCCGCCGGGGTCCGTGCTGTAGTCCCC |
| *4EBP1(MT*-T36A/T46A) | Forward | GGGGACTACGCCGCGGCACCCGGCGGCGCGCTCTTCGCCGCAGCACCGGGAGGTGCCAGG |
| | Reverse | CCTGGCACCTCCCGGTGCTGCGGCGAAGAGCGCGCCGCCGGGTGCCGCGGCGTAGTCCCC |
| *4EBP1(MT*-T37D/T46D) | Forward | GGGGACTACGCCGCGGACCCCGGCGGCGCGCTCTTCGCCGCAGACCCGGGAGGTGCCAG |
| | Reverse | CTGGCACCTCCCGGGTCTGCGGCGAAGAGCGCGCCGCCGGGGTCCGCGGCGTAGTCCCC |
| *4EBP1(MT*-S65A/T70A) | Forward | ATGGAGTGTCGGAACGCCCCTGTGGCCAAAGCACCCCCAAGGGATCTG |
| | Reverse | CAGATCCCTTGGGGGTGCTTTGGCCACAGGGGCGTTCCGACACTCCAT |

| | | |
|---|---|---|
| *4EBP1(MT-*S65D/T70D) | Forward | ATGGAGTGTCGGAACGATCCTGTGGCCAAAGACCCCCCAAGGGATCTG |
| | Reverse | CAGATCCCTTGGGGGGTCTTTGGCCACAGGATCGTTCCGACACTCCAT |
| *4EBP1(YLAA)* | Forward | CGGGAGGTACCAGGATCATCGCTGACCGGAAATTCGCGATGGAGTGTCGGAACTCACC |
| | Reverse | GGTGAGTTCCGACACTCCATCGCGAATTTCCGGTCAGCGATGATCCTGGTACCTCCCG |
| *4EBP1(AAAA)* | Forward | GCTGCAGCCAGACCCCAAGCGCGGCCGCCGCCGCCACTCGCCGGGTGGTGC |
| | Reverse | GCACCACCCGGCGAGTGGCGGCGGCGGCCGCGCTTGGGGTCTGGCTGCAGC |
| *NLS-4EBP1* | Forward-F1 | CAAAGAAGGCTGGACAGGCTAAGAAGAAGAAATACCCTTATGATGTGCCAGA |
| | Forward-F2 | GGTTGCTAGCGCCACCATGAAGCGACCTGCCGCCACAAAGAAGGCTGGACAGGCT |
| | Reverse | GGTTACCGGTGTGCCTCCACCGCTGCCAGATCCGCCAATGTCCATCTCAAACTGTG |
| *NLS-eIF4E* | Forward-F1 | AAAGAAGGCTGGACAGGCTAAGAAGAAGAAAGACTACAAGGACGATGACGA |
| | Forward-F2 | GGTTAAGCTTAAGCGACCTGCCGCCACAAAGAAGGCTGGACAGGCTA |
| | Reverse | GGTTGTCGACAACAACAAACCTATTTTTAG |
| **Primers used in genomic PCR and PCR product sequencing** | | |
| **Gene** | **Primer** | **Sequence (5'-3')** |
| *DEPDC5* allele PCR | Forward | GGCTGCACAGGGCAAGTATG |
| | Reverse | GGGGAAATCAGGCTAGGAACA |
| *CASTOR1* allele PCR | 5’ out | TGCCCTTACCTTGATGGAGC |
| | 3’ out | CCCAAGTCCAAGTGTCAGGG |
| *SAMTOR* allele PCR | 5’ out | GTGTGGTGAAGAGCGTCCAC |
| | 3’ out | TGGCACTATGAATTATACACAGCA |
| | 3’ in | GGGACGTGCAGATCCTTTACATTT |
| *CASTOR1* KI | Forward | AGGAGGTAGTCTCAGCTGGG |
| | Reverse | GTACACAGGTGTCCGCGTAA |
